# Dynamic Bayesian networks for neural information flow: evaluation of continuous and discrete scoring metrics

**DOI:** 10.64898/2026.03.03.709276

**Authors:** Jacob Thomas-Hegarty, Stefan R. Pulver, V Anne Smith

**Author notes:** Contributing authors.

## Abstract

Neural information flow describes the movement of activity between neurons or brain areas. Advances in experimental methods have allowed production of large amounts of observational data related to neuronal activity from the single-neuron to population level. Most current methods for analysing these data are based on pairwise comparison of activity, and fall short of reliably extracting neural information flow network structure. Dynamic Bayesian networks may overcome some of these limitations. Here we evaluate the performance of a range of Bayesian network scoring metrics against the performance of multivariate Granger causality and LASSO regression for their ability to learn the connectivity underlying simulated single-neuron and neuronal population data. We find that discrete dynamic Bayesian networks are the best performing method for single-neuron data, and perform consistently for neural-population data. Continuous dynamic Bayesian networks have a tenancy to learn overly dense structures for both data types, but may have utility in scoping studies on single-neuron data. Multivariate Granger causality is the most robust method for learning structure of neural information flow between neural-populations, but performs poorly on single-neuron data. Significance testing within multivariate Granger causality produces variable results between data types. Overall, this work highlights how the analysis of neural information flow can vary depending on they type and structure of underlying data, and promotes discrete dynamic Bayesian networks as a useful and consistent tool for neural information flow analysis.

## 1 Introduction

Neural information flow (NIF) refers to the flow of activity across brain structures in different behavioural and cognitive states. Understanding how NIF structures change during different behaviours could improve our understanding of how underlying circuits function. For example, in the ex vivo *Drosophila* larval ventral nerve chord, distinct network activity modes are associated with different motor behaviour (Pulver et al., 2015). Understanding the differences in information flow between these network modes may aid in understanding of how these behaviours are brought about by the same anatomical structures. Similarly, pathology linked changes in functional connectivity have been observed in Alzheimer’s disease patients (Yu, Sporns, & Saykin, 2021).

Understanding how this relates to changes in NIF network structures may reveal functional changes that help target novel interventions and improve diagnostics.

Although it constrains NIF, structural connectivity cannot necessarily be used to infer NIF network structure, with some of the strongest structural connections not necessarily providing the strongest pathways of information flow (Benozzo et al., 2024). NIF analysis requires simultaneous observation of neuronal activity in different regions. Such data can be collected using techniques with varying invasiveness and temporal and spatial resolution, such as functional magnetic resonance imaging (fMRI), electroencephalography (EEG), calcium imaging and microarray electrophysiology. Analyses aiming to learn connectivity structures from these types of data may fall into the category of either functional or effective connectivity. Functional connectivity describes statistical associations between activity patterns in brain areas. Effective connectivity attempts to uncover the causal interactions between these units, and generally requires testing of an *a priori* causal neuronal structure, making structure learning difficult (Valdes-Sosa, Roebroeck, Daunizeau, & Friston, 2011). However, likely due to misunderstandings surrounding causal influence, many methods are mislabelled as producing effective connectivity even when they do not uncover causal interactions. As such, these labels have limited utility without understanding a method’s individual assumptions and aims. NIF structures must represent directional and direct connectivity between brain regions. In other words, the network structure must define the direction of flow of influence between regions, and not include any direct connections between regions whose connectivity is mediated by another observed region. This means that, although not necessarily representing causal structures, NIF presents a more thorough approach to understanding activity patterns than many functional connectivity methods, perhaps providing candidate models for future causal analysis.

Currently, it is common to analyse neuronal activity in a pairwise manner, comparing lagged or un-lagged activity of each combination of brain areas, and deeming them connected if this similarity is significant (Wang, Li, Li, & Huang, 2021). These methods fall short of reliable inference of NIF structures for a number of reasons. All un-lagged methods, such as correlation analyses, are unable to infer direction of connectivity between brain regions. Some methods utilise temporal information to infer the direction of influence between a pair of brain areas. Granger causality (GC), and its frequency domain counterpart, partially directed coherence (Baccalá & Sameshima, 2001), are commonly used examples of such methods (Seth, Barrett, & Barnett, 2015; Shojaie & Fox, 2022). Another commonly used method, transfer entropy, is a model free version of the classical GC framework. For Gaussian variables, transfer entropy and GC are equivalent (Barnett, Barrett, & Seth, 2009). Built upon the intuition that a cause must precede its effects, GC quantifies whether the past of a potential parent variable improves prediction on the future of a potential child variable, given the past of the child. However, formation of network structures from individual pairwise comparison often leads to mis-identification of spurious connections between brain regions in the cases of indirect connectivity, common innervation or common sensory input (Novelli & Lizier, 2021). Multivariate Granger causality (MVGC) attempts to allay these concerns through quantification of the improvement in prediction of the future activity pattern of one variable when information on the past of another is included conditioned on the past of all other variables in the system. However, MVGC may still lead to the learning of spurious connections, for example in the case of collider structures in the data generating process or confounding by unobserved variables (Eichler, 2005).

A further limitation of pairwise methods is the arbitrary nature of choosing a threshold on some metric whereby a connection is deemed present in the network, complicating interpretation of results. Significance testing on these metrics may be employed; however, in large networks stringent corrections for multiple testing must be performed, potentially reducing statistical power and increasing occurrence of false negatives (Higgins, Kundu, Choi, Mayberg, & Guo, 2019). Analytical methodologies which learn structures of NIF at the network level, as opposed to interaction level, may provide increased reliability and interpretability of findings. Such methods would be capable of linking entire NIF network structures to behaviour without the need for qualitative threshold selection.

A Bayesian network (BN) is a graphical representation and set of parameters describing the dependence and conditional independence relationships, represented as edges, between a group of variables, represented as nodes. Directed links from a parent to child node represent the statistical dependence of the child on the parent. Each node is conditionally independent of all its non-descendants given its parent nodes. Therefore a BN can map unmediated information flow between its nodes. BN structures can be learned at the network level, without utilising pairwise comparison of node’s activity. These models have been utilised widely in a neuroscience context, particularly in fMRI data (Bielza & Larraeñaga, 2014). Here they have uncovered alterations in NIF associated with mild cognitive impairment (R. Li et al., 2013) and breast cancer treatment (Phillips et al., 2022). However, BN structures may have mathematically equivalent counterparts where the direction of some links are reversed, making interpretation of directionality in such structures problematic. Further, BNs are unable to represent cyclic structures, which is problematic in a NIF context due to the abundance of reciprocal connections within nervous systems (Edelman & Gally, 2013; Lin et al., 2024). These issues limit the utility of static BNs for NIF structure learning.

Through inclusion of temporal information, dynamic BNs (DBNs) overcome these limitations. DBNs have shown strong performance relative to current methods in learning ground-truth connectivity structure in simulated fMRI (Liu, Ji, Yao, & Zhang, 2019; J.C. Rajapakse & Zhou, 2007) and spiking neuronal data, particularly when connections are simulated using highly non-linear synaptic integration (Eldawlatly, Zhou, Jin, & Oweiss, 2008, 2010). NIF structures have been learned as DBNs from a range of data modalities and research areas. Utilising fMRI data, DBN representations of NIF have been used to study changes associated with dementia (Burge, Lane, Link, Qiu, & Clark, 2007; Liu et al., 2019), Parkinson’s disease (J. Li, Wang, & McKeown, 2006), mild cognitive impairment (Ramakrishna & Ramasangu, 2019) schizophrenia (Kim et al., 2008), addiction (L. Zhang, Samaras, Alia-Klein, Volkow, & Goldstein, 2005), ageing (Dang, Chaudhury, Lall, & Roy, 2018) and various cognitive and behavioural states (J. Rajapakse, Wang, Zheng, & Zhou, 2008; J.C. Rajapakse & Zhou, 2007; Warnick et al., 2018; Wu, Wen, Li, & Yao, 2013). Similar methods have been used with EEG (Mutlu & Aviyente, 2009), electrocorticography (H. Zhang et al., 2010), and local field potential data (Smith, Yu, Smulders, Hartemink, & Jarvis, 2006; Tsukahara et al., 2022; Xing et al., 2025) to study NIF related to schizophrenia, hand movement, social gaze, seizures, and auditory circuits.

There are a variety of methodologies for learning the structure of DBNs. Constraint-based methods use conditional independence tests to learn network structure. Alternatively, score based methods search heuristically for the optimal network, utilising scores which aim to quantify the probability of a given DBN structure given the observed data. Score based methods are more widely utilised and may be more accurate (Scutari, Graafland, & Gutiérrez, 2019). There are a variety of scores for learning network structures from either discrete or continuous Gaussian data. Although the former is most commonly used, information lost through discretisation and the summative nature of neuronal influence may mean that continuous scores are advantageous on neuronal data. There is also evidence that they may be more robust to noise on simulated fMRI data than their discrete counterpart (Wu et al., 2013).

The structure of data across spatial scales in neuroscience differs dramatically, from highly non-linear single-neuron dynamics, to more linear oscillatory neuronal population data. Understanding how performance of NIF structure learning methods differ dependent on this context is vital when selecting appropriate analysis tools for a given data type. This may be particularly relevant for continuous DBNs given their assumption of linearity in data streams, but the impacts of these violations are unknown.

Despite some examples of their use in the literature, large scale evaluation of the comparative performance of DBN scores for NIF structure learning from different data types has not been performed, leaving uncertainty around their relative performance with different modalities. This study provides an initial evaluation of range of score based metrics for continuous and discrete DBN structure learning in terms of their ability to reconstruct the connectivity of simulated single-neuron and neural-population data. For comparison, MVGC was also tested, as a frequently used method capable of learning NIF structures in a pairwise manner. Additionally, an unlagged regularized linear regression based method is included for a reference of the performance of unlagged multivariate methods.

## 2 Methods

### 2.1 Simulated datasets

In order to evaluate the performance of structure learning methods, data with known underlying structure is required. Here we utilise two simulated datasets of 100 pseudo-randomly generated 10 node structures. The first dataset was produced using leaky integrate and fire equations, modelling single-neuron dynamics. The second was produced using a neural mass model, modelling the dynamics of populations of neurons. All network generation and simulation was performed in Python (Version 3.13.12).

#### 2.1.1 Single-neuron simulation

Leaky integrate and fire (LIF) equations were used to model single-neuron membrane potentials. Change in sub-threshold membrane potential (*dV*_*k*_) was governed by integration of synaptic inputs with exponential decay towards the resting potential *V*_rest_ at rate *τ*_*m*_:

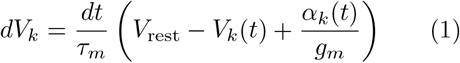

Here *V*_*k*_(*t*) is the membrane potential of neuron *k* at time-point *t, α*_*k*_(*t*) is the total synaptic input to neuron *k* at time-point *t, g*_*m*_ is a fixed parameter describing the resting membrane conductance, and *dt* is the simulation time-step. When a neuron’s membrane potential (*V*_*k*_) reached a threshold (*V*_*th*_), a spike was initiated, and the refractory counter *R*_*k*_ set to *τ*_ref_. At the time-point immediately following this, *V*_*k*_ was set to a firing potential *V*_spike_, representing the depolarisation phase of an action potential. For the following time up to the refractory period *τ*_ref_, *V*_*k*_ was set to *V*_rest_ :

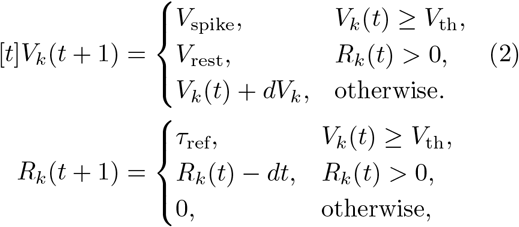

Networks were formed of 10 neurons. Each neuron sent four connections to different randomly selected neurons, with uniform probabilities of any non-self connection being present. Three neurons were randomly selected to be inhibitory. A total of 100 structures were simulated.

Synaptic input *α* to each neuron *k* at time-step *t* was modelled using conductance based synapses:

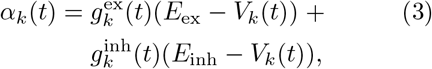

Where 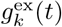 and 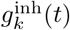 are the excitatory and inhibitory synaptic conductances of neuron *k* at time-point *t*. Intuitively, when 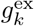 or 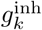 are high, membrane potential in the postsynaptic neuron *k* will tend towards the relevant synaptic equilibrium potentials *E*_Ex_ and *E*_inh_. Conductances increase by Δ*g*_ex_ and Δ*g*_inh_ when the neuron receives a spike from a connected excitatory or inhibitory neuron. Otherwise they decay exponentially, with the rate of this decay controlled by the synaptic time parameters *τ*_ex_ and *τ*_inh_:

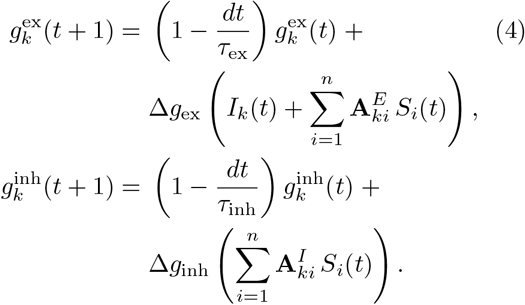

**A**^*E*^ and **A**^*I*^ represent excitatory and inhibitory connectivity matrices. *n* is the number of neurons in the network, so that 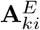 and 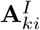 indicate the presence or absence of an excitatory or inhibitory connection from neuron *i* to neuron *k. S*_*i*_(*t*) is 1 if neuron *i* fired at time-point *t* and 0 otherwise. The summation terms count the number of connected excitatory or inhibition neurons which fired at time-point *t*. Exogenous spikes are represented by *I*_*k*_(*t*), which is 1 if there is an exogenous input spike to *k* at time *t* and 0 otherwise. Each neuron received exogenous stimulation via an independent Poisson spike train with a firing rate of 25Hz, generated via sampling of inter-spike intervals from an exponential distribution with a mean of 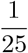, modelling input to the network from unobserved excitatory neurons.

Parameter values related to neuronal dynamics were selected based on those used in previously in the literature (Vogels & Abbott, 2005). Synaptic variables Δ*g*_ex_ and Δ*g*_inh_ were selected based on a parameter search, optimising these values to maximise the proportion of random networks with stable firing rates and irregular firing patterns.

The latter was measured through calculation of the coefficient of variation of inter-spike intervals (CV_ISI_). The selected values and interpretation of fixed parameters in the above equations are shown in Table 1.

**Table 1.**
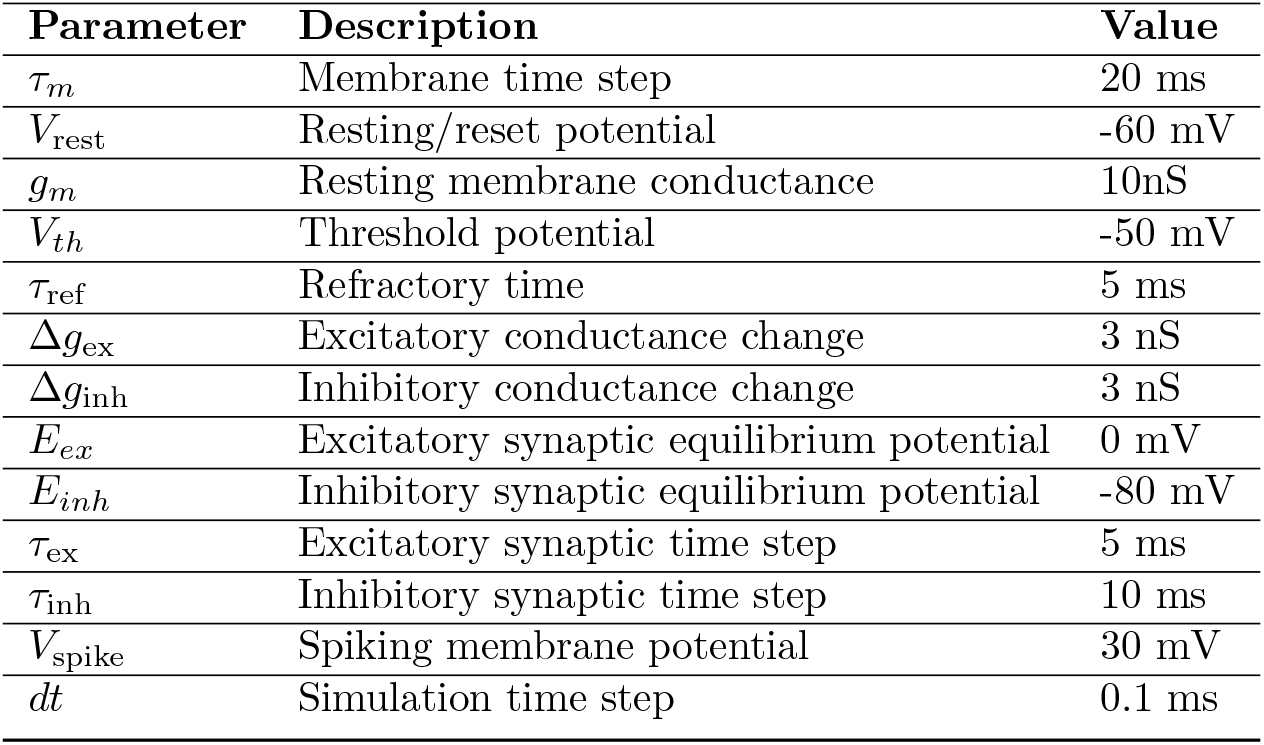
Parameters utilised in leaky integrate and fire simulations.

| Parameter | Description | Value |
| --- | --- | --- |
| $\tau_m$ | Membrane time step | 20 ms |
| $V_{\text{rest}}$ | Resting/reset potential | -60 mV |
| $g_m$ | Resting membrane conductance | 10nS |
| $V_{th}$ | Threshold potential | -50 mV |
| $\tau_{\text{ref}}$ | Refractory time | 5 ms |
| $\Delta g_{\text{ex}}$ | Excitatory conductance change | 3 nS |
| $\Delta g_{\text{inh}}$ | Inhibitory conductance change | 3 nS |
| $E_{\text{ex}}$ | Excitatory synaptic equilibrium potential | 0 mV |
| $E_{\text{inh}}$ | Inhibitory synaptic equilibrium potential | -80 mV |
| $\tau_{\text{ex}}$ | Excitatory synaptic time step | 5 ms |
| $\tau_{\text{inh}}$ | Inhibitory synaptic time step | 10 ms |
| $V_{\text{spike}}$ | Spiking membrane potential | 30 mV |
| $dt$ | Simulation time step | 0.1 ms |

Simulations were run for 100 s, with a time-step (*dt*) of 0.1 ms. Data was averaged into 1 ms bins to provide 100,000 data-points for the structure learning process.

#### 2.1.2 Neural-population simulation

Wilson–Cowan oscillators are a form of neural mass model which separately model the activity of excitatory and inhibitory neurons within neuronal populations (Wilson & Cowan, 1972). These activities are generally described by two partial differential equations modelling all possible interactions between these units (Figure 2):

**Fig. 1.**
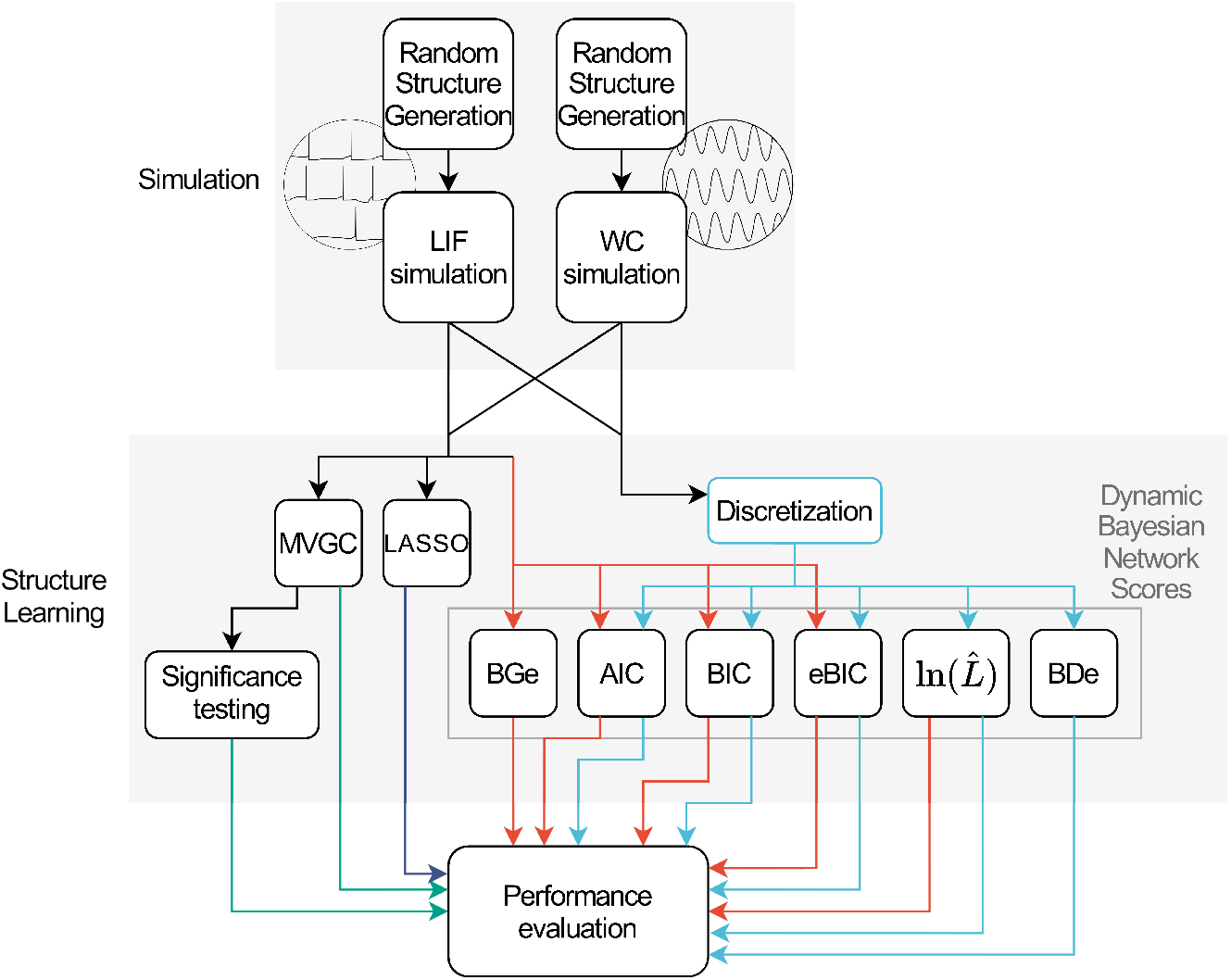
Study workflow. Two sets of randomly generated connectivity structures were generated. Activity across one set was simulated using leaky integrate and fire dynamics, and the other using a Wilson-Cowan based model of connected oscillators. Data from these simulations were used to test a range of dynamic Bayesian network learning scores, multivariate Granger causality (MVGC) and LASSO regression based methods for their ability to re-learn the structures underlying the simulated data. Where possible, the methods were tested on both discrete and continuous data. (BGe: Bayesian Gaussian equivalent, AIC: Akaike information criterion, BIC: Bayesian information criterion, eBIC: extended Bayesian information criterion, ln(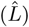) : log-likelihood, BDe: Bayesian Dirichlet equivalent)

**Fig. 2.**
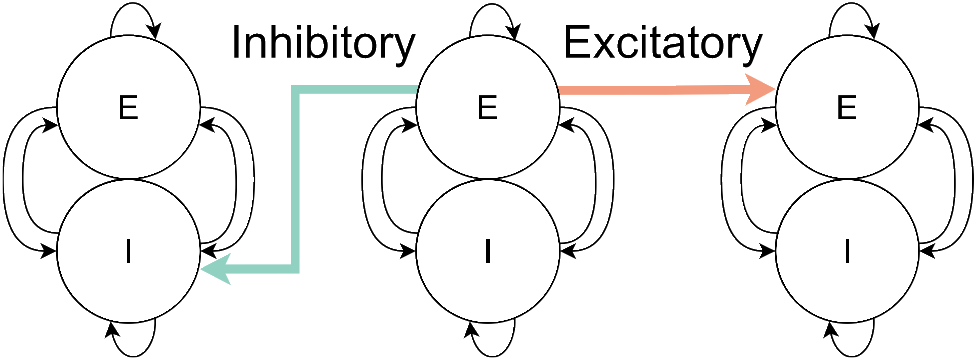
The structure of neural-populations and their exogenous connections used in neural-population simulations. One unit represents the basic structure of a Wilson Cowan oscillator, illustrating the connectivity between subpopulations. Connections between units illustrate the inter-population connectivity used to randomly connect populations in neural-population simulations. Excitatory connections between populations are represented by connecting respective excitatory subpopulations (red arrow), and inhibitory connections by connecting the excitatory subpopulation of the parent oscillator to the inhibitory subpopulation of the child oscillator (green arrow).

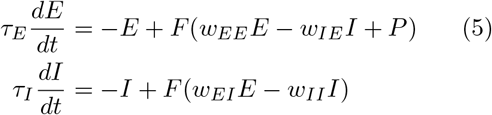

where *E* and *I* represent excitatory and inhibitory subpopulation activity, *τ*_*E*_ and *τ*_*I*_ represent the excitatory and inhibitory population time-steps, and *P* represents the exogenous excitatory input to the population. The *w* terms represent connectivity between subpopulations, with *wEI* representing the connectivity strength from the inhibitory to the excitatory subpopulation, *wIE* representing the connectivity strength from the excitatory to the inhibitory subpopulation, and *wEE* and *wII* representing the strength of endogenous connectivity within these sub-populations. In this work, the activation function *F* was a shifted sigmoidal function with threshold of *θ* and gain of *α*:

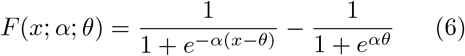

The basic Wilson-Cowan equations were adapted to allow simulation of distinct connected neuronal populations with stochastic exogenous input in discrete time:

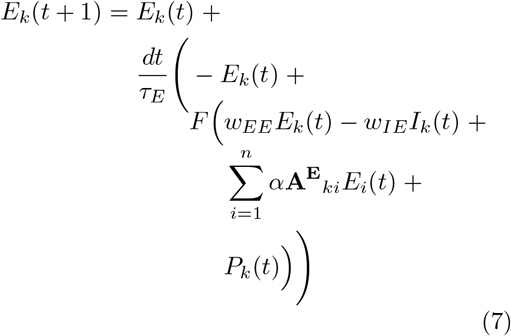

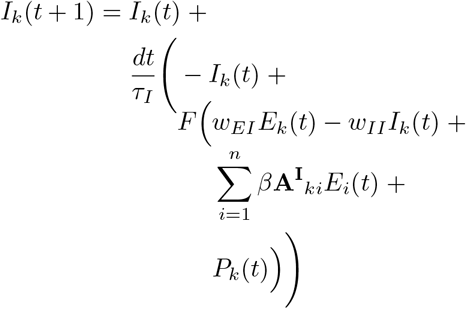

where *E*_*k*_ and *I*_*k*_ are the excitatory and inhibitory sub-population activity of population *k*, **A**^*E*^ and **A**^*I*^ are excitatory and inhibitory connectivity matrices, and *α* and *β* the excitatory and inhibitory inter-population connectivity weights. *n* is the number of neurons in the network, so that 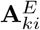 and 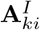 indicate the presence or absence of a connection from the excitatory sub-population of *i* to the excitatory or inhibitory population of *k* (Figure 2), and the summation terms calculate the weighted input activity from inter-population connections. *P*_*k*_(*t*) is the exogenous input noise to population *k* at time-step *t*, modelled as independent Orstein-Uhlenbeck (OU) processes:

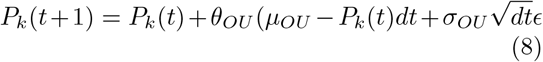

where *ϵ* is a random variable sampled from a normal distribution of mean 0 and variance 1. *µ*_OU_, *θ*_OU_, and *σ*_OU_ are fixed parameters controlling the mean, rate of reversion to the mean, and volatility of the OU process respectively. The values of all fixed parameters were selected to reliably exhibit chaotic activity patterns, as measured using the coefficient of variation of inter-peak intervals (*CV*_*IP I*_) across randomly connected 10 neuron networks, and are displayed in Table 2.

**Table 2.**
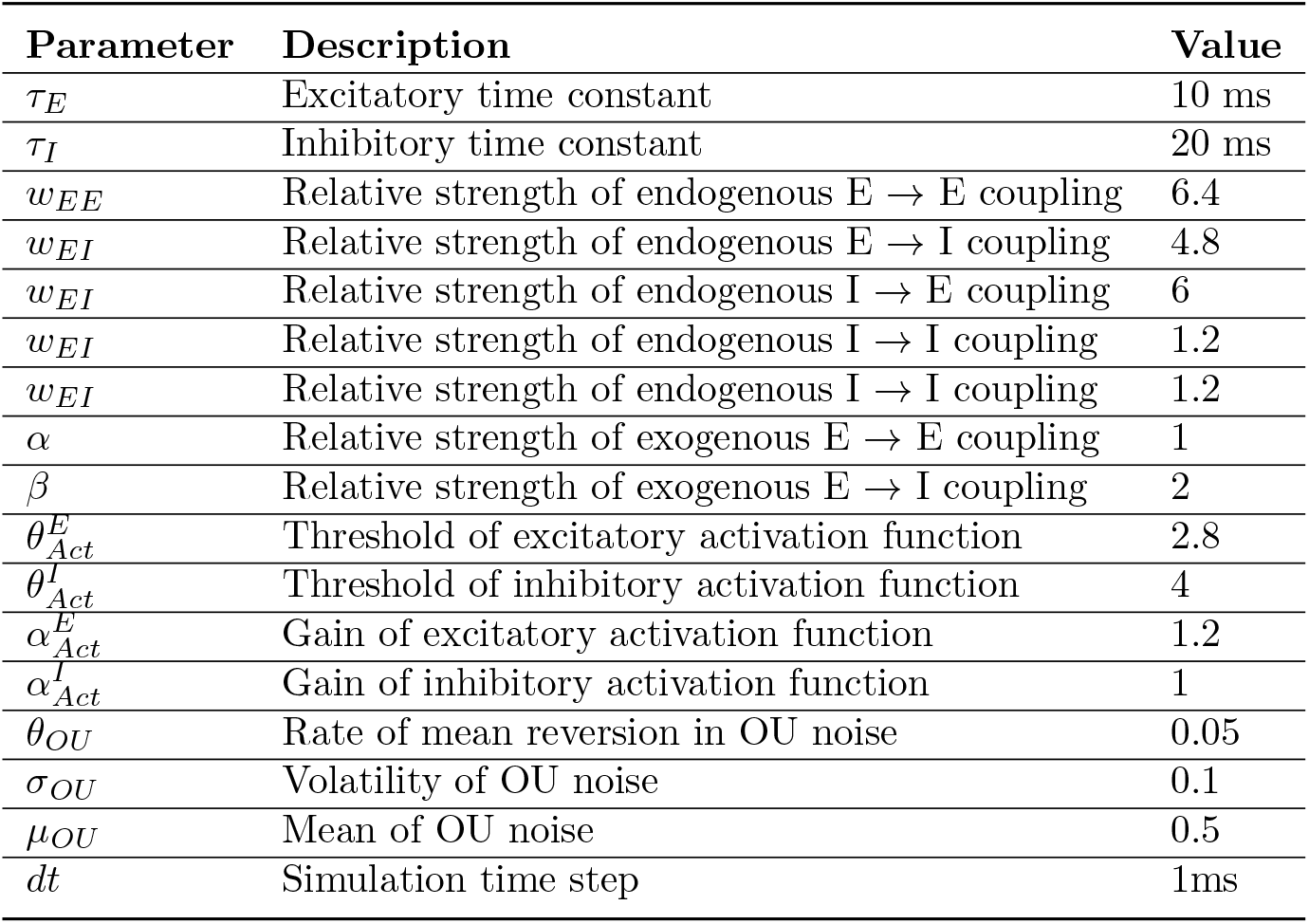
Parameters utilised in Wilson-Cowan simulations.

| Parameter | Description | Value |
| --- | --- | --- |
| $\tau_E$ | Excitatory time constant | 10 ms |
| $\tau_I$ | Inhibitory time constant | 20 ms |
| $w_{EE}$ | Relative strength of endogenous E → E coupling | 6.4 |
| $w_{EI}$ | Relative strength of endogenous E → I coupling | 4.8 |
| $w_{EI}$ | Relative strength of endogenous I → E coupling | 6 |
| $w_{EI}$ | Relative strength of endogenous I → I coupling | 1.2 |
| $w_{EI}$ | Relative strength of endogenous I → I coupling | 1.2 |
| $\alpha$ | Relative strength of exogenous E → E coupling | 1 |
| $\beta$ | Relative strength of exogenous E → I coupling | 2 |
| $\theta_{Act}^E$ | Threshold of excitatory activation function | 2.8 |
| $\theta_{Act}^I$ | Threshold of inhibitory activation function | 4 |
| $\alpha_{Act}^E$ | Gain of excitatory activation function | 1.2 |
| $\alpha_{Act}^I$ | Gain of inhibitory activation function | 1 |
| $\theta_{OU}$ | Rate of mean reversion in OU noise | 0.05 |
| $\sigma_{OU}$ | Volatility of OU noise | 0.1 |
| $\mu_{OU}$ | Mean of OU noise | 0.5 |
| $dt$ | Simulation time step | 1ms |

Inter-population connectivity matrices were formed in a similar manner to the LIF simulations, although here 30% of connections were randomly selected as inhibitory, meaning a population may send both inhibitory and excitatory inter-population connections. Excitatory connections resulted in the excitatory unit of the parent population influencing the excitatory unit of the child population (Figure 2). Inhibitory connections resulted in the excitatory unit of the parent influencing the inhibitory unit of the child population. Simulations were run for 1000 s, with a time-step (*dt*) of 1 ms. Data was averaged into 10 ms bins to provide 100,000 data-points for the structure learning process.

#### 2.1.3 Simulation metrics

*CV*_*ISI*_ and *CV*_*IP I*_ values were calculated as the ratio of the standard-deviation to the mean of the inter-spike and inter-peak intervals, with a value close to 1 indicating irregular firing pattens. A cross-correlation analysis was used to estimate the true lag between connected nodes in the simulated networks. In this analysis, only nodes which were directly connected in simulations were tested. The lag at which the correlation between the parent and child was maximised was recorded, and the distribution of these values across all connections and simulations were considered for each simulation type.

### 2.2 Learning network structure

#### 2.2.1 Dynamic Bayesian networks

Dynamic Bayesian networks (DBNs) extend the principle of BNs across the temporal domain. In a DBN a node at a given time-point is dependent on itself at each previous time-point from the minimum to the maximum Markov lag, as well as parent nodes at any of these time points (Smith, 2010). The graphical representation of these nodes across time can then be reduced down to a single network, representing the cross-node dependencies in the dynamic network. The resultant network is able to overcome the key limitations of a static BN. Firstly, this network may contain cycles, as arcs may represent reciprocal relationships between nodes at different time points. Additionally, it is possible to confidently infer the direction of an arc, as causal influence must be exerted forwards in time. The maximum and minimum Markov lag in all discussed analyses is 1. To force the necessary connections within the DBN structure, forward links from a variable to the lagged version of itself are required to be present. This means that self-connections are always present and therefore are not considered in performance evaluation. Links backwards in time are prevented, asserting the assumption that influence moves forward in time. DBN structures were handled using the DBayesNet package, developed alongside this work to allow DBN structure learning in R with bnlearn (Scutari, 2010) and available at https://github.com/jacobhegarty/DBayesNet.

DBN structures may be learned in the same manner as static BNs. There are a range of score-based metrics available to evaluate network structures, a variety of which were evaluated here. These scores all evaluate the network structure as a whole for its ability to describe the data through calculating the probability of the candidate graphical structure given the observed data. Using Bayes theorem, with *G* representing the candidate graphical structure and *D* the observed data, this can be calculated as:

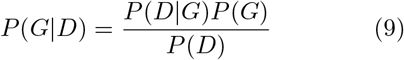

For the purposes of structure learning, the probability of observing the data *P* (*D*) is constant and thus ignored. In practice, the probability of the candidate graphical structure graphical structure *P* (*G*) is often kept constant, although it can be utilised to include prior information over the likelihood of different graph structures. This leaves the probability of observing the data given it was generated by the candidate graphical structure *P* (*D*|*G*) to be calculated. This can be done using scores based on maximum or marginal likelihood, many of which have formulations for both continuous and discrete data. For discrete scores, quantile discretisation was performed to label data into three classes using the Hmisc library in R (version 5.2.3) (Harrell Jr, 2025). All scoring metrics were applied using bnlearn (Version 5.0.2) (Scutari, 2010). See section 2.3 for a brief overview of the scoring metrics tested here.

The number of possible network structures increases super-exponentially with the number of network nodes, and finding the best scoring network is an NP-hard problem (Chickering, 1996). This means that scoring of all networks to find the optimal solution is not possible, and heuristic search strategies must be utilised. Here, searches were performed in bnlearn using a hill-climb algorithm (Scutari, 2010), a method which has been widely employed in the literature (McNally, Mair, Mugno, & Riemann, 2017; Owens et al., 2022; Smith et al., 2006; Tsukahara et al., 2022). This strategy is a type of greedy search which starts with a random network structure and randomly adds, deletes, or reverses the direction of a link. If this action improves the network score, it is accepted into the network structure. This process is repeated until no improvements to the network score can be made in this manner (Smith, 2010). This solution is not guaranteed to represent the overall best scoring network, as the search may have become stuck on a local minima, where network score cannot be improved by making a single change but does not represent the best network overall. In order to negate this risk, the hill-climb search was repeated 100 times, each beginning with a different random structure. Initial investigations indicated that this number of repeats was sufficient to find the same scores as longer searches, indicating that the best scoring network had been found.

#### 2.2.2 Multivariate Granger causality

Conceptually, GC tests whether information on the past of variable *X* improves prediction on the future of variable *Y* beyond the past of *Y* itself (Granger, 1969). In practice, this involves fitting a ‘reduced’ and ‘full’ model, the former being an autoregressive model for the dynamics of *Y* up to a maximum lag *L* and the latter a vector autoregressive (VAR) model of *Y* as a combination of the past of *Y* and *X* up to lag *L*. The residuals of these models can then be compared to calculate the Granger causality from *X* to *Y*. In the multivariate case (MVGC), the GC of *X* on *Y* is conditioned on other variables in the system, with these additional variables being included in both the full and reduced models (Geweke, 1984). Granger causality is computed as

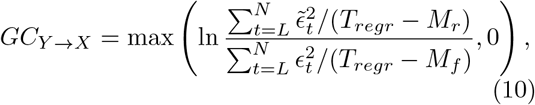

where *N* is the number of time-points, *L* is the model order or lag, and 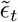 and *ϵ*_*t*_ are the residuals from the reduced and full models at time t. This means that MVGC is calculated as the logged ratio of the sum of squared residuals from the reduced 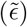 and full (*ϵ*) models, normalized by the difference between the number of time points used in the models (*T*_*regr*_ = *T* − *L*) and the number of parameters in the corresponding reduced and full regression models *M*_*r*_ and *M*_*f*_ (where *M*_*r*_ = *M*_*f*_ − *L*).

Selection of model order *L* is important to balance the fit of the VAR model. Here, the elbow point of the average Akaike Information Criterion (AIC) value for all models in the network as a function of lag was used. Given that the underlying dynamics do not change between simulations, and computational cost of testing performing multiple lagged regressions for each potential connection across all datasets, this subanalysis was limited to 10 simulations, with the value of *L* used on the whole dataset selected as the point at which increasing the *L* produced negligible change in average AIC for all simulations tested.

The significance of MVGC values can be utilised via comparing residuals 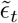 and *ϵ*_*t*_ using an F-test. Two types of Granger causality networks were formed, “significant-only” with links selected based on significant GC values, and “non-zero” with links selected for all non-zero MVGC values. A 0.05 significance threshold was adjusted for the 90 potential connections within the network using a Bonferroni correction, meaning the criteria for a link in the significant-only MVGC networks was 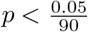.

#### 2.2.3 LASSO regression

A LASSO regularisation forces the absolute values of regression coefficients to be less than the parameter *λ*, shrinking the coefficients of less important variables towards zero. In order to learn network structure, each node was modelled using all other neurons, with all nodes with non-zero coefficients selected as its parents. Optimal *λ* values were selected for each regression as the largest value which gives a model error within 1 standard error of the minimum. This method selects the simplest (least parametrised) model with accuracy within a reasonable distance of the most accurate model, thereby balancing avoidance of over-fitting and model accuracy (Friedman, Hastie, & Tibshirani, 2010a). LASSO regressions and *λ* selection were performed using the glmnet package in R (Version 4.1.8) (Friedman, Hastie, & Tibshirani, 2010b).

### 2.3 Bayesian network scoring metrics

#### 2.3.1 Maximum likelihood scores

A number of scoring metrics are based on estimating the probability of the data given the graph and the maximum likelihood estimation of the parameters (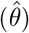) from the data (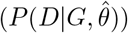).

##### Log-likelihood

The simplest of these is the log-likelihood, the logged value of the maximum likelihood (Koller & Friedman, 2009). For discrete data this is calculated as:

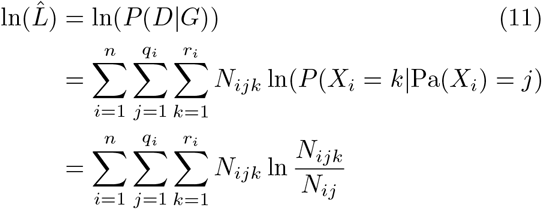

where *n* is the number of nodes, *q*_*i*_ is the number of parent set configurations for node *X*_*i*_, *r*_*i*_ is the number of possible states of node *X*_*i*_, *N*_*ijk*_ is the number of instances where the parents of *X*_*i*_ (Pa(*X*_*i*_)) are in configuration *j* and *X*_*i*_ is in state *k*, and *N*_*ij*_ is the number of times a the parent set of *X*_*i*_ are in state 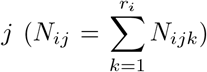. Therefore, the discrete log-likelihood is the sum of the logged probability of each child state given each parent state, weighted by the number of times this combination occurs in the data.

For continuous data, assuming a joint Gaussian distribution with independent identically distributed observations, the log-likelihood score may be calculated as

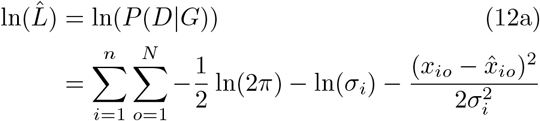

where *x*_*io*_ is the observed value of child node *X*_*i*_ at observation *o*, 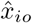 is the fitted value of linear regression of the parent variables on the child variable *i* 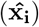 at observation *o*.

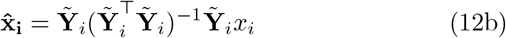

and *σ*_*i*_ is the standard deviation of residuals of this regression. 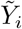 is all observations of *Pa*(*X*_*i*_) and *x*_*i*_ is all observations of *X*_*i*_.

Intuitively, log-likelihood will find the structure in which the parents contain the maximum possible information on the child. As such, this score favours complex structures, often learning fully connected networks, and is included here as a reference point. The reliance on linear regression in the continuous log-likelihood score means that this score and its penalised versions (below) are only able to capture linear relationships between nodes.

##### Akaike information criterion (AIC)

The Akaike information criterion (Akaike, 1998):

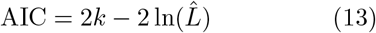

extends this score to include a penalty term proportional to the number of parameters in the model (*k*). This penalty attempts to limit model complexity, with structures containing more connections being more strongly penalised.

##### Bayesian information criterion (BIC)

The Bayesian information criterion (Schwarz, 1978):

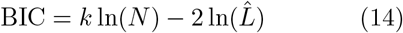

penalises ln 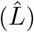 in proportion to both *k* and the number of data points used in structure learning (*N*). The penalty term in BIC is more severe than that of AIC when *N >* 7, which is generally the case in BN structure learning.

##### Extended Bayesian information criterion (eBIC)

The extended Bayesian information criterion (Foygel & Drton, 2010):

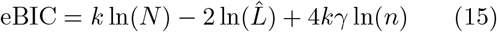

score adds an additional penalty relative to both *k* and the size of the network (*n*). The parameter *γ* controls the severity of this penalty, and the default value of 0.5 was used here. In the structure learning process, the size of the network remains equal so this the additional penalty has the effect of scaling complexity penalisation by network size. For *γ* = 0.5 and a network of 10 nodes, eBIC applies a more severe complexity penalty of 2 ln(10)*k* relative to BIC.

#### 2.3.2 Marginal likelihood scores

As opposed to the aforementioned maximum likelihood scores, the Bayesian Dirichlet equivalent (BDe) and Bayesian Gaussian equivalent (BGe) scores do not rely on maximum likelihood estimation of parameters, or contain an explicit penalty term for complexity. They instead calculate the marginal likelihood of the data given the candidate graphical structure (*P* (*D* | *G*)) by integrating over the likelihood of the data given all possible values for parameters *θ* for this structure, weighted by a prior over the value of these parameters:

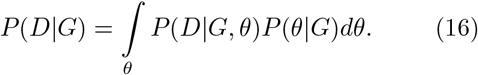

Bayesian Dirchichlet (BD) scores (Heckerman, Geiger, & Chickering, 1995) are utilised for discrete data, and contain a Dirichlet prior (the conjugate prior of the multinomial distribution) for the distribution of probabilities for each state occurring in a node given its parents’ states, for all possible combinations and configurations of parents. This results in a network being scored as:

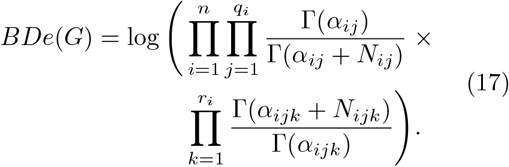

Where *n* represents the number of nodes in the network, *q*_*i*_ the number of parent configurations for node *X*_*i*_, and *r*_*i*_ the number of states of node *X*_*i*_. The hyper-parameter *α*_*ijk*_ controls the Dirichlet distribution, and 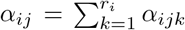. As in the discrete log-likelihood score *N*_*ijk*_ is the number of times each child-parent configuration combination is observed in the data, 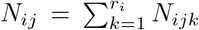. Intuitively, the second product is higher when the parents of each node best predict the corresponding child, and the first product is higher when there are fewer parents for each node and with an equal number of observations of each parent configuration. The first term, therefore, provides an implicit penalty for the complexity of the structure. For BDe the parameters of the Dirichlet prior are calculated from the the imaginary sample size (iss) parameter *α*, which was set to 1 for all analysis in this study, via the equation: 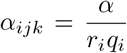. Utilising a uniform prior removes the need to subjectively specify a prior for each nodes’ potential parent configurations and assumes that all BN structures are equally likely. In reference to this, BDe may also be referred to as BDeu in the literature, throughout this paper we refer to BDeu as BDe. It is worth noting that BDe is closely related to another popular BN scoring algorithm K2, which uses *α*_*ijk*_ = 1 (Cooper & Herskovits, 1991).

The BGe score (Geiger & Heckerman, 1994) is used for continuous data, utilising a Normal-Wishart distribution (the conjugate prior of a multivariate Gaussian distribution) for the mean and precision matrix parameters of the joint normal distribution over each parent-child node set. This results in a network being scored as:

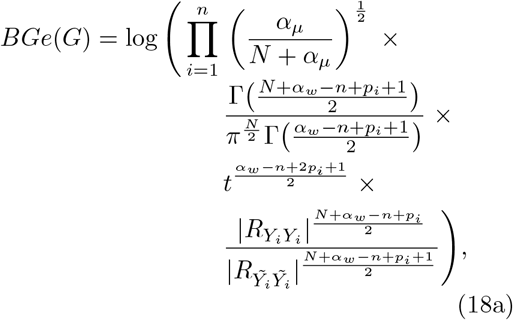

Where

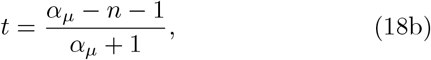

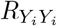 is the posterior scale matrix of parent and child nodes,

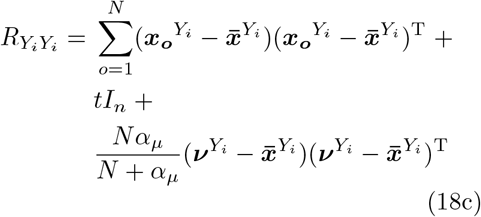

and 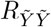 is the same posterior scale matrix without the child node *X*_*i*_:

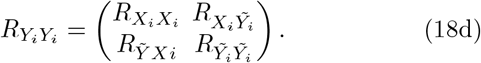

As in BDe, *N* is the total number of observations, and *n* the total number of nodes, in the data. *p*_*i*_ is the number of parents of node *X*_*i*_ in the candidate structure. Set *Y*_*i*_ contains the node *X*_*i*_ and its parents in the candidate structure, and 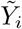 contains only the parents of node *X*_*i*_. 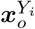 is the vector of observations of all nodes in *Y*_*i*_ at observation *o*. 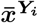 is a vector containing the mean values of nodes in *Y*_*i*_, and 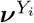 is the vector containing the priors over node means for nodes in Y. In this work, the prior over node means is calculated as the mean value of that node, meaning the last term in the calculation of 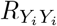 disappears. 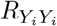 is therefore the un-normalised covariance, or scatter matrix, of *Y*_*i*_, with *t* added to the diagonal values of this matrix corresponding to the variance of each variable in *Y*_*i*_ via addition of the product of *t* and an identity matrix of size n (*I*_*n*_). 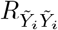 is calculated in the same manner, but for set 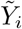. The final term in the BGe calculation therefore measures how much adding the child to its parents increases the total covariance. If the child is well explained by its parents, this increase is small, and the score is higher. The previous terms come from integration over parameters and the Normal-Wishart prior, and include an implicit penalty for structure complexity and lack of observations. *α*_*w*_ and *α*_*µ*_ are the equivalent sample size hyper-parameters controlling this prior over the spread of the precision matrix and mean respectively. These were set to *n* + 2 and 1 for all work presented here. As with the Gaussian log-likelihood based scores, data are assumed to be sampled from a joint Gaussian distribution, with independent and identically distributed observations. This results in the score only accounting for linear relationships between variables.

### 2.4 Performance evaluation

Performance was evaluated using precision and recall. Precision measures the ability of a method to find true links without finding false links (Equation 19).

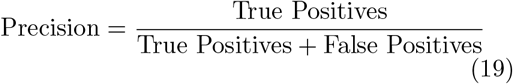

Recall measures the ability of the method to find true links (Equation 20). High precision and low recall indicate under-fitting, whereas low precision and high recall indicate over-fitting.

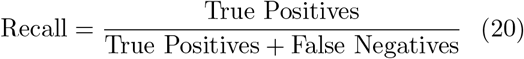

It is important to consider both of these statistics together as a high score in one but not the other cannot indicate high performance. High recall with low precision is characteristic of over-fitting, and low recall with high-precision is characteristic of under-fitting in network structures.

Hypergeometric testing was also performed to ascertain the probability of achieving the same number of true positives as a method, given the number of arcs predicted in the learned network, if picking at random. This was performed separately for each score on each simulation, providing a distribution of performance, whereby a low probability indicates the method performed better than chance.

## 3 Results

### 3.1 Single-neuron data

Figure 3 shows a one second example LIF simulation. The mean *CV*_*ISI*_ of LIF simulations was 0.93 (*±* 0.19 across simulations), with an average within simulation standard deviation of 0.20 (*±* 0.11 across simulations). The mean firing rate was 35.77 Hz (*±* 21.51 Hz across simulations), with a mean within simulation variation of 26.32 Hz (*±* 14.67 Hz across simulations). This indicates that the simulated structures produced a large range of network dynamics. A cross-correlation analysis found a median and mode optimal lag of 0 between all connections present in every simulation.

**Fig. 3.**
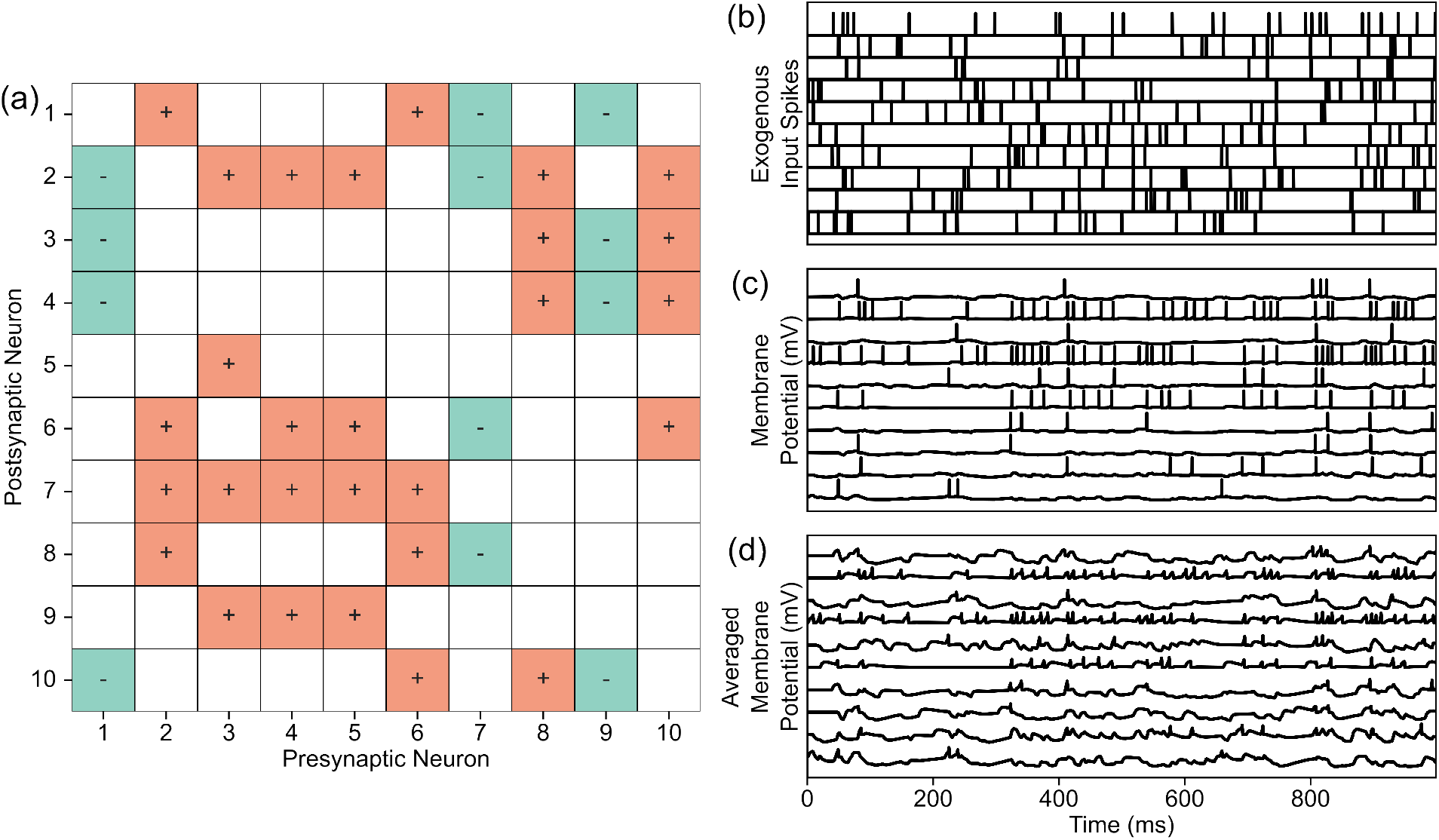
Example figures from a single leaky integrate and fire simulation run. (a) Example leaky integrate and fire network structure. Red squares indicate excitatory connections and green squares indicate inhibitory connections. (b) Example exogenous input to leaky integrate and fire networks. (c) Example simulated membrane potential trace made using network structure and input shown in a and b. (d) Simulation in c averaged in 1ms bins, as used in structure learning process.

The average performance of all tested structure learning methods on the single-neuron data in precision-recall space is shown in figure 4. Both log-likelihood scores completely over-fit these data, with high recall (both 1.00) and low precision (both 0.44) (Figure 4). This is due to them learning fully connected network structures 100% of the time with discrete data, and 93% of the time with continuous data.

**Fig. 4.**
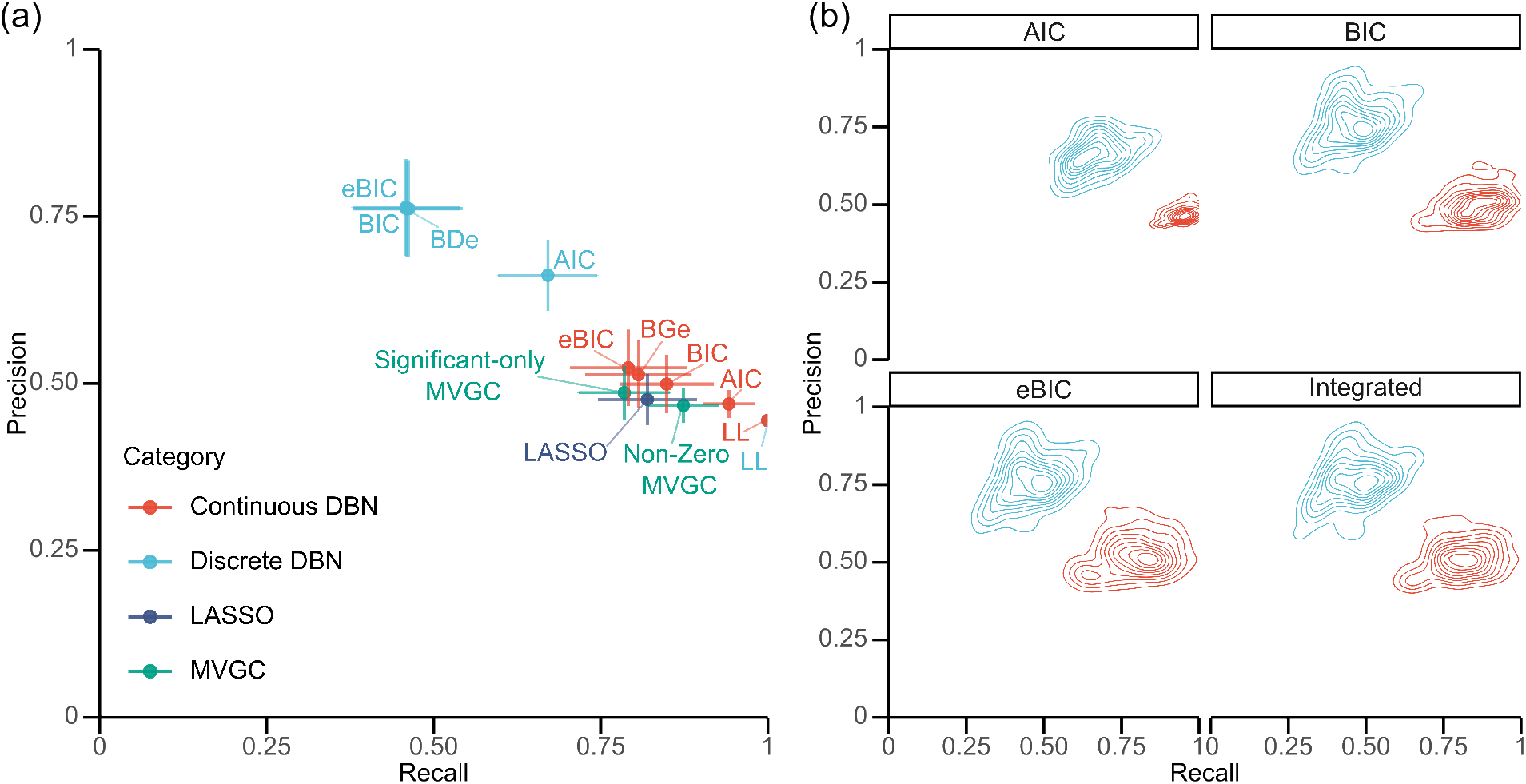
a) The average performance of discrete (blue) and continuous (red) DBN scores (eBIC = extended Bayesian information criterion, BIC = Bayesian information criterion, AIC = Akaike information criterion, LL = log-likelihood, BDe = Bayesian Dirichlet equivalent, BGe = Bayesian Gaussian equivalent), LASSO (purple) and MVGC (green) across 100 single-neuron leaky integrate and fire simulations when compared to ground truth connectivity, plotted in precision-recall space with recall on X-axis and precision on Y-axis. b) The distribution in performance of DBN score types on discrete and continuous data for 100 single-neuron simulations, when compared to ground truth connectivity. Produced using a 2-D kernel density estimate calculated independently for each group with 10 bins separated by contour lines representing increasing density.

Mean precision and recall for discrete BIC, eBIC and BDe cluster closely in both precision and recall, all with an average precision of 0.76 and recall of 0.46 (Figure 4). This indicates an underfit of this dataset. Discrete AIC achieves a more balanced performance, with an decreased average precision (0.66) and increased average recall (0.67) (Figure 4). Interestingly, discrete AIC predicts closest to the true number of connections in the network, with an average of 40.60, compared to the ground truth of 40. All other discrete scores, other than log-likelihood, underestimate the number of connections present (Figure 5). Hypergeometric testing indicated a low average probability of achieving equal or better performance by chance for all discrete scores other than log-likelihood (all mean p*<*0.01; Figure 5).

**Fig. 5.**
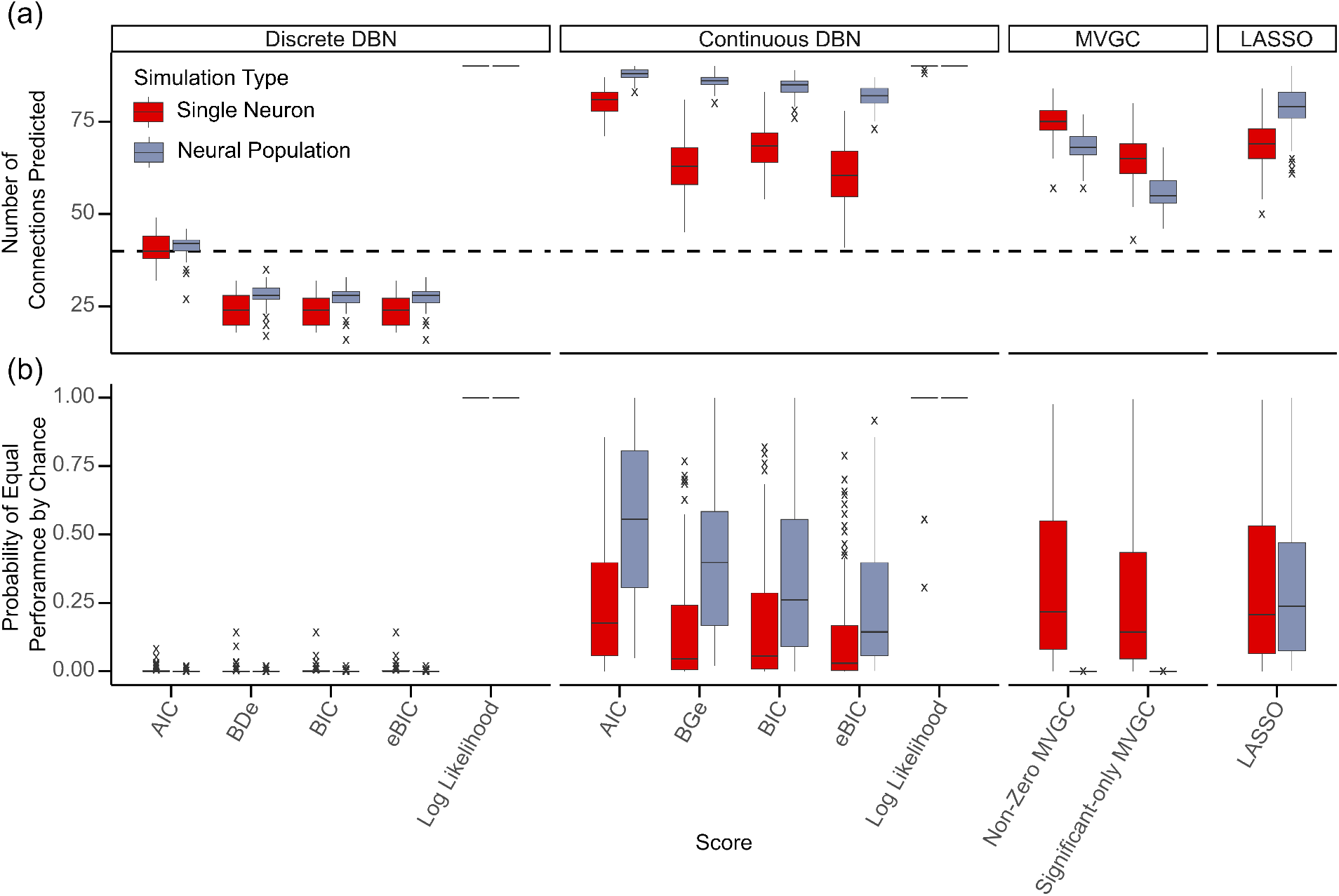
a) Number of connections predicted to be in structure underlying simulated single-neuron (red, N=100) and neural-population data (purple, N=100). Dashed line indicates the true number of connections in each network (40). b) Probability of predicting the same or more correct connections as each structure learning method, given the number of connections they predicted, from simulated single-neuron (red, N=100) and neural-population data (purple, N=100). Calculated using the hypergeometric distribution. Central hinge represents median value. Lower and upper hinges correspond to first and third quartiles. Whiskers indicate the furthest point within a 1.5 interquartile range from the hinge. Crosses indicate outliers.

Continuous DBNs sit much further to the right in precision-recall space, with higher recall and lower precision. There is more separation in the performance of these score’s than their discrete counterparts, with increasing recall and decreasing precision from eBIC (precision = 0.52, recall = 0.79), to BGe (precision = 0.51, recall = 0.81), to BIC (precision = 0.50, recall = 0.85), to AIC (precision = 0.47, recall = 0.94) (Figure 4). The continuous scores tended to overestimate the number of connections in the network (Figure 5). Hypergeometric testing indicated a moderate probability of achieving equal or better performance by chance for the continuous scores other than log-likelihood, with AIC having the highest probability at 0.26 with a large standard deviation of 0.25, and eBIC the lowest at 0.13 with a standard deviation of 0.20. (Figure 5). Aside from log-likelihood, there is a clear separation in precision-recall space between the continuous and discrete version of each score with continuous scores over-fitting and discrete scores under-fitting data (Figure 4).

Optimal model order for MVGC was estimated as 10 (Figure 8). Average recall for significant-only MVGC networks was 0.79, with an average precision of 0.49. For non-zero GC networks, average recall increased to 0.87 and average precision decreased to 0.47. Viewed in precision-recall space, it is clear that these methods perform extremely poorly on the single-neuron simulations (Figure 4). Both significant-only and non-zero MVGC tended to over-estimate network density, with 64.87 and 74.91 connections respectively (Figure 5). Hypergeometric testing further indicates high probabilities of achieving these levels of performance by chance, with an average of 0.33 for non-zero and 0.25 for significant-only networks. (Figure 5).

The non-lagged LASSO method had an average recall of 0.82 and precision of 0.48 on the LIF dataset (Figure 4). This method tended to over-estimate the number of connections, with an average of 69.07 (Figure 5), indicating a clear over-fit of this dataset. The average probability of observing the same performance by chance was 0.31 (Figure 5).

### 3.2 Neural-population data

Figure 6 shows a 10 second example WC simulation. Simulations displayed patterns of chaotic oscillatory activity. A cross-correlation analysis comparing connected excitatory populations in all networks gave a distribution of optimal lags with a median of 1 and a mode of 2.

**Fig. 6.**
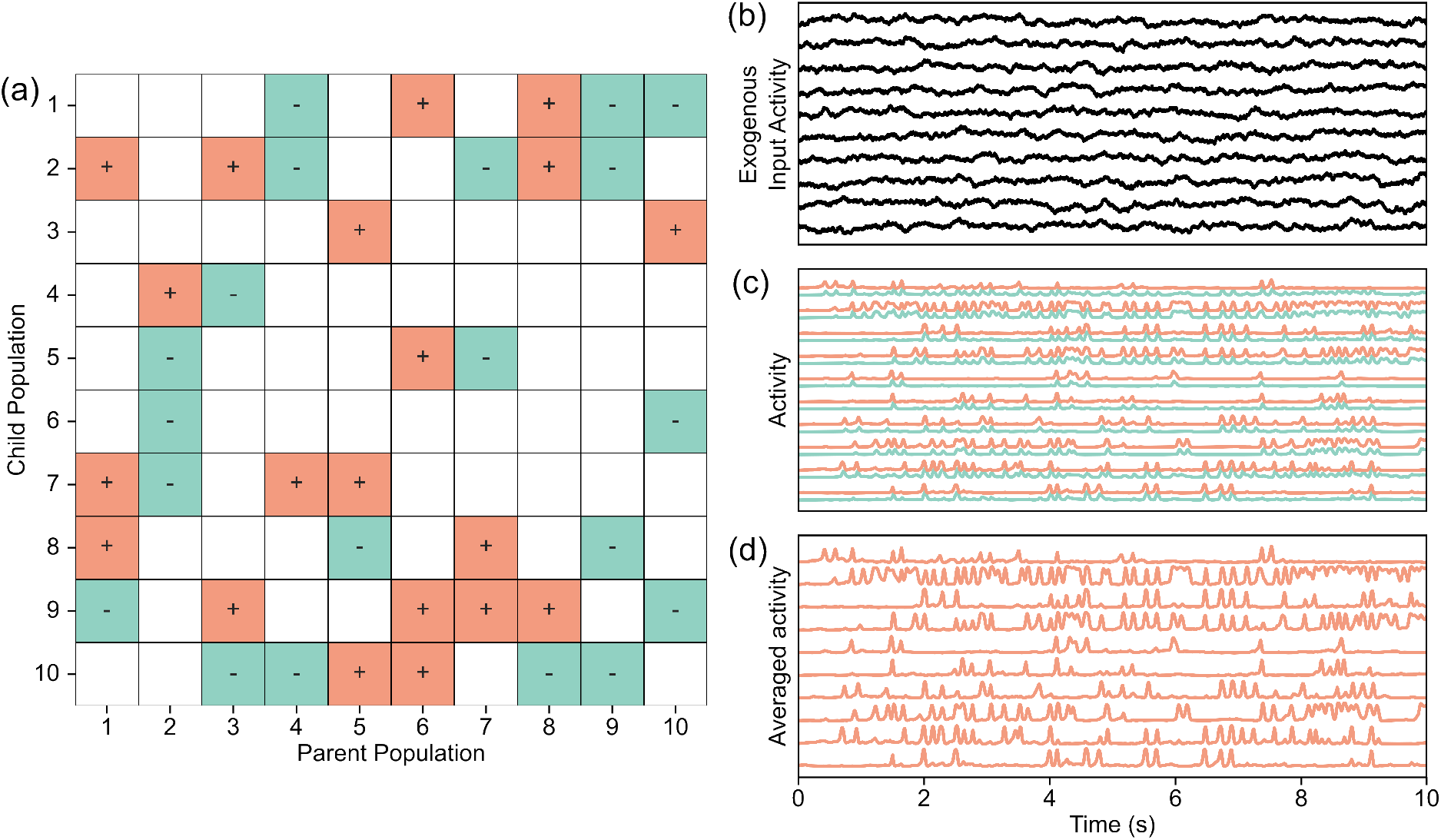
Example figures from a single Wilson-Cowan simulation run. (a) Example Wilson-Cowan network structure. Red squares indicate excitatory connections and green squares indicate inhibitory connections. (b) Example exogenous input to Wilson-Cowan network. (c) Example simulated population activity trace made using network structure and input shown in a and b. Excitatory unit activity shown in green, and inhibitory unit activity shown in red. Excitatory activity is offset vertically to aid visualisation. d) Excitatory trace averaged in 10ms bins, as used in structure learning process.

The mean *CV*_*IPI*_ of excitatory neurons in neural-population simulations was 0.60 (*±* 0.08 across simulations), with an average within simulation standard deviation of 0.17 (*±* 0.04 across simulations). The mean number of peaks in per second was 4.68 (*±*0.533 across simulations) excitatory traces, with a mean within simulation variation of 1.38 (*±* 0.443 across simulations). The mean *CV*_*IPI*_ of inhibitory neurons in neural-population simulations was 0.55 (*±*0.09 across simulations), with an average within simulation standard deviation of 0.16 (*±*0.07 across simulations). The mean number of peaks per second in inhibitory sub-populations was 5.11 (*±*0.557 across simulations) in inhibitory traces, with a mean within simulation variation of 1.08 (*±*0.446 across simulations).

The average performance of all tested structure learning methods on the neural-population data is shown in figure 7. As with the single-neuron data both the discrete and continuous log-likelihood scores over-fit data, both learning fully-connected network structures for every neural-population simulation (Figure 5).

**Fig. 7.**
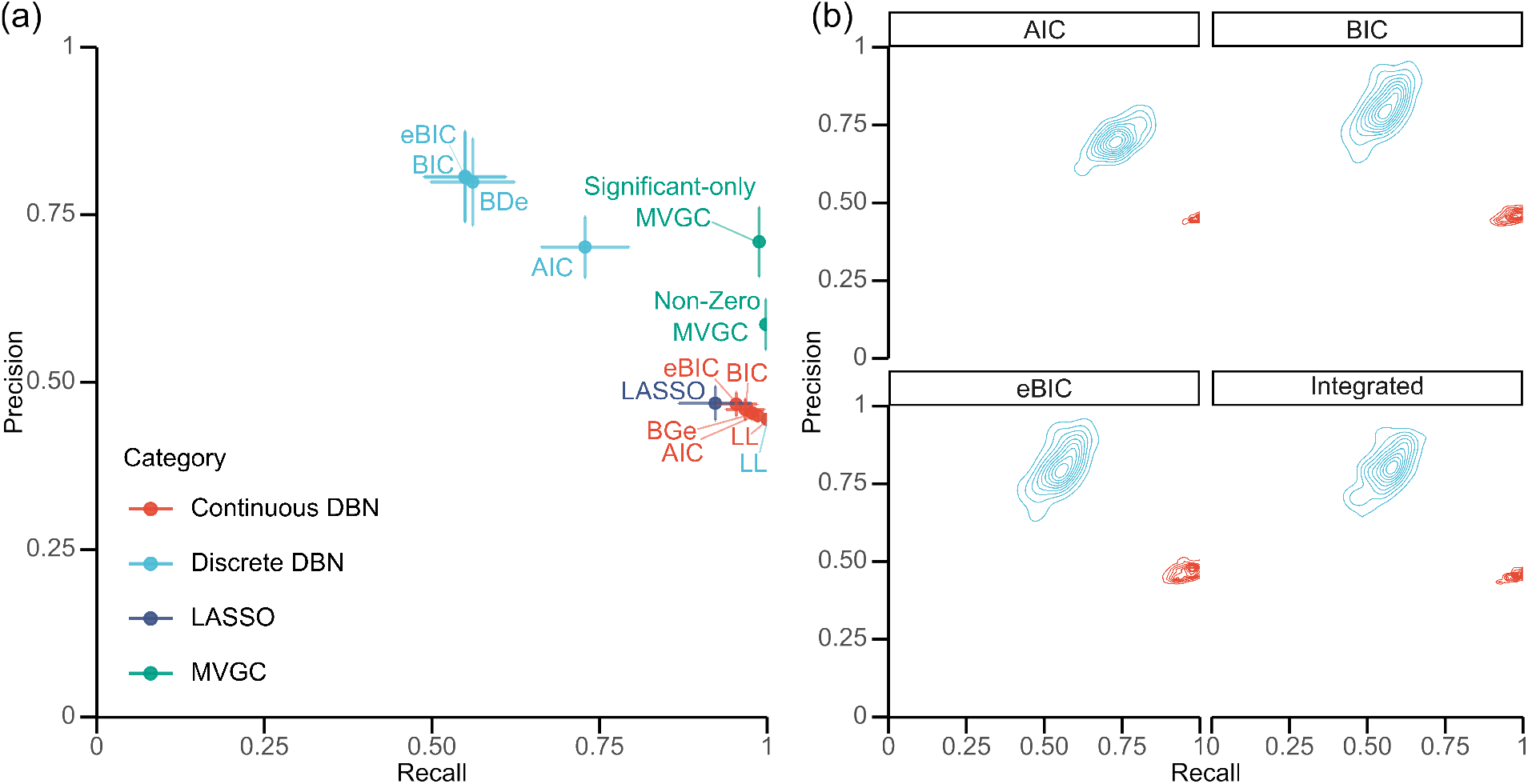
a) The average performance of discrete (blue) and continuous (red) DBN scores (eBIC = extended Bayesian information criterion, BIC = Bayesian information criterion, AIC = Akaike information criterion, LL = log-likelihood, BDe = Bayesian Dirichlet equivalent, BGe = Bayesian Gaussian equivalent), LASSO (purple) and MVGC (green) across 100 neural-population Wilson-Cowan simulations, when compared to ground truth connectivity. b) The distribution in performance of DBN score types on discrete and continuous data for 100 neural-population simulations, when compared to ground truth connectivity. Produced using a 2-D kernel density estimate calculated independently for each group with 10 bins separated by contour lines representing increasing density.

Discrete e-BIC, BIC and BDe scores cluster closely in precision-recall space, eBIC and BIC with a recall of 0.55 and precision of 0.81 and BDe with a recall of 0.56 and precision of 0.80 (Figure 7). As with the single-neuron data, discrete AIC was the best performing BN scoring metric for neural-population data, with an average precision of 0.70 and recall of 0.73 (Figure 7). Discrete AIC predicted 41.50 connections on average, and all other discrete scores other than log-likelihood under-estimated network density (Figure 5). Hypergeometric testing indicated that all discrete scores other than log-likelihood had a low average probability of being equalled or outperformed by chance (all mean p*<*0.001; Figure 5).

Continuous scores all appear to over-fit neural-population data, with very high recall and low precision (AIC precision = 0.45, recall = 0.99; e-BIC precision = 0.47, recall = 0.95; BIC precision = 0.46, recall = 0.97; BGe precision = 0.45, recall = 0.98) (Figure 7). Continuous scores over-estimated network density, all predicting over 80 connection on average (Figure 5), with AIC learning fully connected structures 7% of time, BGe 1% and all other scores 0%. The average probability of observing the same performance as these methods by chance was 0.54 for AIC, 0.41 for BGe, 0.34 for BIC and 0.25 for e-BIC (Figure 5).

Optimal model order for MVGC analysis was estimated as 7 (Figure 8). Significant-only MVGC achieved an average precision of 0.71 and average recall of 0.99. Non-zero MVGC had a higher average recall of 1, but lower average precision of 0.59 (Figure 7). Both significant-only and non-zero MVGC tended to over-predict network density, with an average of 55.98 and 68.33 predicted connections respectively (Figure 5). Hypergeometric testing indicated an extremely low average probability of either of these methods being equalled or outperformed by chance (both mean p*<*0.0001; Figure 5).

**Fig. 8.**
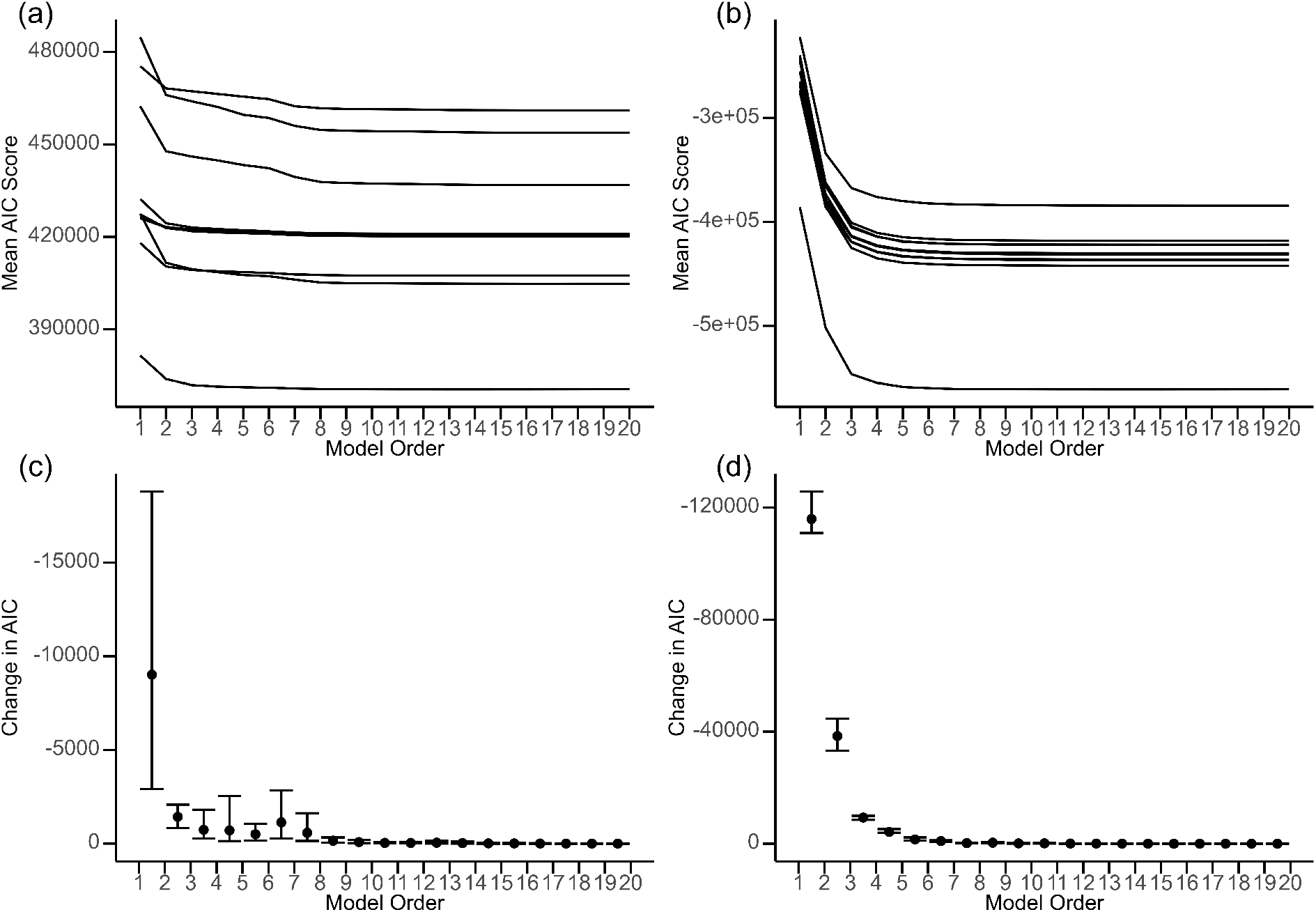
The upper panel shows average AIC values for full model of every node in 10 a) single-neuron and b) neural-population simulations with increasing model order, with each line representing a single simulation. The lower panel shows change in average AIC with increasing model order for the same 10 c) single-neuron and d) neural-population simulations, with error bars representing the range across the 10 simulations tested. Optimal model order for each data type was selected as the minimum value past which increasing order produces negligible change in AIC.

The non-lagged LASSO method had a mean recall of 0.92, and close-to-chance average precision of 0.47 (Figure 7). This method tended to overestimate network density, with 78.96 connections predicted on average (Figure 5) and 1% of learned networks fully connected. The average probability of achieving the same or better performance as LASSO by chance was 0.30 (Figure 5).

## 4 Discussion

We have evaluated the relative performance of discrete and continuous DBNs, MVGC and unlagged LASSO regressions in learning the structures underlying simulated single-neuron and neural-population data. We have found continuous DBNs to over-fit both data types, with a large number of false positive connections being learned, to the extent of performing close to chance on the neural-population data. Discrete DBNs achieve a more balanced performance on both datasets, although have a tenancy to produce false negatives. MVGC performs poorly on single-neuron data, but well on neural-population data, especially when significance testing is utilised. Unlagged LASSO regression performs extremely poorly, strongly over-fitting both of these data types.

Both the single-neuron and neural-population datasets include a wide range of network dynamics. Large between-sample variance in mean spiking rate and frequency indicate a range of excitability in network structures. Large between-sample variance in the CV for spikes and peaks indicates a range in regularity of activity across the datasets. This variance improves the generalisability of our findings to applications across nervous systems, and was the main justification for utilising random, as opposed to more stable or predefined biological network structures.

The dynamics underlying the single-neuron data were much quicker than the data resolution utilised in structure learning, with cross-correlation analysis of the links in ground truth structures indicating a maximal correlation in unlagged data. This may explain the poor performance of all methods on these data. With an extremely large number of observations, structure learning becomes slower, presenting the challenging problem of balancing the length of recordings with temporal resolution. It may be possible to increase temporal resolution whilst maintaining recording length when learning a single structure, as opposed to the hundreds of repeated learning tasks here. However, the findings of Eldawlatly *et al*. Eldawlatly et al. (2010) suggest that reducing the Markov lag bellow the true synaptic lag causes a much more severe reduction in performance than observed here. In the neural-population model, cross-correlation analysis of existing connections gave a median optimal lag of 1ms and a mode of 2ms. The comparability of these values to the resolution of neural-population data used in structure learning likely contributed to the increased performance of discrete DBNs and MVGC compared to the single-neuron data. Interestingly, there does not appear to be a benefit to continuous DBNs, which over-fit neural-population much more severely than single-neuron data.

Where they separate in performance, the penalised log-likelihood scores follow a clear pattern of increasing connection density of learned networks from e-BIC to BIC to AIC. This reflects relaxation of the complexity penalty through this sequence of scores, from 2 ln(10)*k* + *k* ln(100000) in e-BIC, to *k* ln(100000) in BIC, to 2*k* in AIC. Interestingly, the positioning of the integrated scores within this trend appears to be dependent on data type and vary between BDe and BGe. BDe appears to cluster closely with e-BIC and BIC on both datasets, slightly outperforming these scores on the neural-population data. The similarity in performance of BIC and BDe, both learning sparser structures than AIC, aligns with previous findings (Ke, Keenan, & Smith, 2025). BGe sits between e-BIC and BIC in precision-recall space on single-neuron data and over-fits slightly more than these scores in the population data. As expected due to their lack of penalty for complexity, the log-likelihood score tends to learn fully connected structures, and is not a useful metric for network structure learning in this context. Although discrete AIC appears to achieve the best balance of precision and recall of all BN scoring metrics on both data types, the most appropriate scoring metric for a given study will be highly contextual. Given the heavy trade-off between precision and recall for all metrics, particularly on the single-neuron data, the researcher must choose the acceptable balance in risk of missing true versus finding spurious connections.

There is a clear separation of discrete and continuous score performance across all BN scores tested and both data types. These results suggest that the increase in information contained in continuous activity traces lead to over-fitting of data. This over-fitting may be attributed to the small noise fluctuations in continuous traces which would not have affected discretised labels. This is in contrast to Wu et al. (2013), who found that Gaussian DBNs learned using the BIC score were more robust to increasing noise than their discrete counterparts. This difference in findings may be due to the increased complexity of simulations utilised here, with Wu *et al*. utilising linear multivariate autoregressive models which may have favoured Gaussian over discrete DBNs. In single-neuron data, continuous networks may offer an advantageous increase in recall for scoping studies where reliability of findings is less important than identifying potential connections. In population level data, however, the over-fitting effect is so extreme that results are unlikely to be useful in any context. This result is surprising given the increased coherence of the population model to the assumption of linearity of continuous DBNs.

Both non-zero and significant-only MVGC networks perform poorly on the single-neuron data, often performing close to chance and worse than all DBN scores other than log-likelihood. In this case, significance testing of links appears to cause a decrease in overall performance, with a moderate increase in average precision accompanied by a large decrease in recall. Clearly, in this case more true positives than false positives were filtered out by significance testing. In contrast, when viewed in precision-recall space it is clear that significant-only MVGC achieves the best performance on neural-population data. In this case non-zero MVGC performs with a much lower average precision, suggesting that here significance testing has successfully filtered out false positive links. The difference in effects of significance testing on MVGC performance between data types illustrates the difficulties in thresholding link selection in pairwise methods, and the ambiguity of proper statistical techniques in such conditions.

The contrast in relative performance of DBNs and MVGC on different data types, with MVGC the best performing on more linear neuronal systems and DBNs on more non-linear systems, provides guidance on the most appropriate methods in these situations. The fact that the most common optimal lag for neural-population data based on cross-correlation analysis was greater than the maximum Markov lag used in DBN structure learning, but less than the model order used in MVGC, may explain the increase performance of the latter method on this data type. Unfortunately, learning DBN structures with a lag compatible to that used by MVGC is generally unfeasible. For example, a model order of 7 in MVGC corresponds to a DBN with a minimum Markov lag of 1 and a maximum Markov lag of 7. In a rolled network of 10 nodes, this corresponds to a dynamic network of 80 nodes and 2^1890^ potential structures, and the computational complexity of searching heuristically across a space this large would make structure learning unfeasible. On single-neuron data, however, DBNs outperform MVGC, illustrating differences in the ability of these methodologies to capture more noisy non-linear neuronal dynamics. This may have been expected due to the reliance of MVGC on linear modelling Dang, Chaudhury, Lall, and Roy (2015), however contrasts with the results of Chen et al. (2023) that MVGC may still be suitable for quantifying interactions in non-linear systems. Unsurprisingly, the un-lagged LASSO method performs poorly on both datasets, highlighting the importance utilising lag-based methods for NIF analysis on any data type.

Given the evidence collected here, the choice of the best structure learning method depends on the researcher’s goals and data. Spiking neuronal data is highly noisy and non-linear, and DBNs appear to deal with these data better than MVGC. For a scoping study, aiming to identify potential connections within a network where precision is not important, continuous DBNs may be preferable. For studies which require increased confidence in the validity of learned structures, discrete DBNs will be more suitable. The discrete AIC achieves the most balanced performance. As more neurons’ spiking activity is collated, for example by methods from calcium imaging to EEG, and data becomes increasingly linear (Deco, Jirsa, Robinson, Breakspear, & Friston, 2008), MVGC with significance testing achieves the best performance. It is never recommendable to use non-lagged methods, such as the LASSO regression represented here.

This study has provided an initial investigation of DBN structure learning performance on neuronal data. These models are meant to be generally biologically plausible, but are not intended to emulate any specific neural network. Despite allowing generalisability to a range of network dynamics, this means that structures do not correspond to a specific biological network. The parameters utilised within simulations were selected to produce stable, irregular dynamics across a range of random structures. This removed biological relevance from some of the parameters utilised, as the processes they represent are likely to vary between circuits *in vivo*. Additionally, the variation in structure may have produced subtle differences in the lag of information flow. This means that estimated optimal model order in MVGC may differ slightly between simulations, and use of the same model order within simulation types may have effected these results. However this effect will be minimal, given that parameters remain constant through simulations.

Given discrete DBNs consistency across data types relative to MVGC, future work may focus on development of these metrics to further improve their performance, providing a versatile and reliable tool for NIF structure learning. Given the number of structures learned in this paper, a maximum Markov lag of 1 was utilised throughout analysis. Future work may focus on quantifying the benefits of increasing this value for data with larger and more variable tue lags, such as the neural-population data utilised here. Additionally, the iss parameter utilised in BDe may be increased to encourage this method to learn an increased number of connections (Steck & Jaakkola, 2002). This may improve the fit of models learned using this score, encouraging BDe to produce networks with increased recall. Previous results suggest this effect may be especially beneficial in large datasets such as those available in neuroscience Ke et al. (2025); Ulusoy and Geduk (2024). It is likely, however that this increase in recall will be accompanied by a decrease in precision, and further work is required to investigate how increasing this parameter changes the positioning of BDe in precision recall space. Another benefit of DBNs comes from the ability to incorporate structural information into the scoring process. This was not tested here due to the nature of the simulation meaning information flow is defined by the structural connectivity. However, there has been some success in combining information from fMRI and DTI in the structure learning process (Dang et al., 2018). This may be especially beneficial for single cell data from organisms such as *Drosophila*, where the structural connectome is well characterised (Azevedo et al., 2024; Dorkenwald et al., 2024; Takemura et al., 2024; Winding et al., 2023).

## 5 Conclusion

This work has shown continuous DBNs to overfit simulated single-neuron and neural-population data, to the extent that they perform close to chance on the latter data type. Discrete DBNs achieved a better balance, although tended to learn overly sparse networks. Discrete AIC score achieved the highest performance of all DBN scores, and the highest performance overall on single-neuron data. MVGC performed close to chance on single-neuron data, but achieved high performance on neural-population data when utilising significance testing over connection presence. This same significance testing depleted performance on single-neuron data. Taken together these findings illustrate the tendency of continuous DBNs to learn much denser structures than their discrete counterparts on neuronal data, the importance of considering data type when selecting structure learning methods, and the ambiguity around correct statistical testing measures when utilising MVGC. Future work may focus on optimising discrete DBNs for use on neuronal data, potentially developing them into a versatile tool across neuronal data types.

## 6 Declarations

### 6.2 Code availability

The code used in this paper is available at: https://github.com/jacobhegarty/DBN-scores-neuro. This includes use of the R package DBayesNet, which was developed alongside this work and is available at https://github.com/jacobhegarty/DBayesNet.

## 7 Appendix

### 7.1 MVGC model order

## References

Akaike, H. (1998). Information theory and an extension of the maximum likelihood principle. In E. Parzen, K. Tanabe, & G. Kitagawa (Eds.), Selected papers of hirotugu akaike (pp. 199–213). New York, NY: Springer New York.

Azevedo, A., Lesser, E., Phelps, J.S., Mark, B., Elabbady, L., Kuroda, S., … Tuthill, J.C. (2024). Connectomic reconstruction of a female drosophila ventral nerve cord. Nature, 631 (8020), 360–368, 10.1038/s41586-024-07389-x

Baccalá, L.A., & Sameshima, K. (2001). Partial directed coherence: A new concept in neural structure determination. Biological Cybernetics, 84 (6), 463–474, 10.1007/pl00007990

Barnett, L., Barrett, A.B., Seth, A.K. (2009). Granger causality and transfer entropy are equivalent for gaussian variables. Physical Review Letters, 103 (23), 238701, 10.1103/physrevlett.103.238701

Benozzo, D., Baron, G., Coletta, L., Chiuso, A., Gozzi, A., Bertoldo, A. (2024). Macroscale coupling between structural and effective connectivity in the mouse brain. Scientific Reports, 14 (1), 3142, 10.1038/s41598-024-51613-7

Bielza, C., & Larraeñaga, P. (2014). Bayesian networks in neuroscience: A survey. Frontiers in Computational Neuroscience, 8, 131, 10.3389/fncom.2014.00131

Burge, J., Lane, T., Link, H., Qiu, S., Clark, V.P. (2007). Discrete dynamic bayesian network analysis of fmri data. Human Brain Mapping, 30 (1), 122–137, 10.1002/hbm.20490

Chen, X., Ginoux, F., Carbo-Tano, M., Mora, T., Walczak, A.M., Wyart, C. (2023). Granger causality analysis for calcium transients in neuronal networks, challenges and improvements. eLife, 12, e81279, 10.7554/eLife.81279

Chickering, D.M. (1996). Learning bayesian networks is np-complete. In D. Fisher & H.J. Lenz (Eds.), Learning from data: Artificial intelligence and statistics v (pp. 121–130). New York, NY: Springer New York.

Cooper, G.F., & Herskovits, E. (1991). A bayesian method for constructing bayesian belief networks from databases. Proceedings of the seventh conference on uncertainty in artificial intelligence (p. 86–94). San Francisco (CA): Morgan Kaufmann.

Dang, S., Chaudhury, S., Lall, B., Roy, P.K. (2015). Assessing assumptions of multi-variate linear regression framework implemented for directionality analysis of fmri. 2015 37th annual international conference of the ieee engineering in medicine and biology society (embc) (p. 2868–2871).

Dang, S., Chaudhury, S., Lall, B., Roy, P.K. (2018). Tractography-based score for learning effective connectivity from multimodal imaging data using dynamic bayesian networks. IEEE Transactions on Biomedical Engineering, 65 (5), 1057–1068, 10.1109/TBME.2017.2738035

Deco, G., Jirsa, V.K., Robinson, P.A., Breakspear, M., Friston, K. (2008). The dynamic brain: From spiking neurons to neural masses and cortical fields. PLoS Computational Biology, 4 (8), 1–35, 10.1371/journal.pcbi.1000092

Dorkenwald, S., Matsliah, A., Sterling, A.R., Schlegel, P., Yu, S.-c., McKellar, C.E., … FlyWire Consortium (2024). Neuronal wiring diagram of an adult brain. Nature, 634 (8032), 124–138, 10.1038/s41586-024-07558-y

Edelman, G.M., & Gally, J.A. (2013). Reentry: A key mechanism for integration of brain function. Frontiers in Integrative Neuroscience, 7,, 10.3389/fnint.2013.00063

Eichler, M. (2005). A graphical approach for evaluating effective connectivity in neural systems. Philosophical Transactions of the Royal Society B: Biological Sciences, 360 (1457), 953–967, 10.1098/rstb.2005.1641

Eldawlatly, S., Zhou, Y., Jin, R., Oweiss, K. (2008). Reconstructing functional neuronal circuits using dynamic bayesian networks. 2008 30th annual international conference of the ieee engineering in medicine and biology society (p. 5531–5534).

Eldawlatly, S., Zhou, Y., Jin, R., Oweiss, K.G. (2010). On the use of dynamic bayesian networks in reconstructing functional neuronal networks from spike train ensembles. Neural Computation, 22 (1), 158–189, 10.1162/neco.2009.11-08-900

Foygel, R., & Drton, M. (2010). Extended bayesian information criteria for gaussian graphical models. Proceedings of the 24th international conference on neural information processing systems - volume 1 (p. 604–612). Red Hook, NY, USA: Curran Associates Inc.

Friedman, J., Hastie, T., Tibshirani, R. (2010a). Regularization paths for generalized linear models via coordinate descent. Journal of Statistical Software, 33 (1), 1–22, 10.18637/jss.v033.i01

Friedman, J., Hastie, T., Tibshirani, R. (2010b). Regularization paths for generalized linear models via coordinate descent. Journal of Statistical Software, 33 (1), 1–22, 10.18637/jss.v033.i01

Geiger, D., & Heckerman, D. (1994). Learning gaussian networks. R.L. de Mantaras & D. Poole (Eds.), Uncertainty in artificial intelligence (p. 235–243). San Francisco (CA): Morgan Kaufmann.

Geweke, J.F. (1984). Measures of conditional linear dependence and feedback between time series. Journal of the American Statistical Association, 79 (388), 907–915, 10.1080/01621459.1984.10477110

Granger, C.W.J. (1969). Investigating causal relations by econometric models and cross-spectral methods. Econometrica, 37 (3), 424–438, 10.2307/1912791

Harrell Jr, F.E. (2025). Hmisc: Harrell miscellaneous [Computer software manual]. (R package version 5.2-5)

Heckerman, D., Geiger, D., Chickering, D.M. (1995, September). Learning bayesian networks: The combination of knowledge and statistical data. Machine Learning, 20 (3), 197–243, 10.1007/BF00994016

Higgins, I.A., Kundu, S., Choi, K.S., Mayberg, H.S., Guo, Y. (2019). A difference degree test for comparing brain networks. Human Brain Mapping, 40 (15), 4518–4536, 10.1002/hbm.24718

Ke, X., Keenan, K., Smith, V.A. (2025). Evaluation of bayesian network scoring functions in polychotomous data analysis. Discover Data, 3 (1),, 10.1007/s44248-025-00033-7

Kim, D., Burge, J., Lane, T., Pearlson, G., Kiehl, K., Calhoun, V. (2008). Hybrid ica–bayesian network approach reveals distinct effective connectivity differences in schizophrenia. NeuroImage, 42 (4), 1560–1568, 10.1016/j.neuroimage.2008.05.065

Koller, D., & Friedman, N. (2009). Probabilistic graphical models: principles and techniques. Cambridge, MA: MIT Press.

Li, J., Wang, Z., McKeown, M. (2006). Dynamic bayesian networks (dbns) demonstrate impaired brain connectivity during performance of simultaneous movements in parkinson’s disease. 3rd IEEE International Symposium on Biomedical Imaging: Macro to Nano, 2006., 964, 10.1109/isbi.2006.1625080

Li, R., Yu, J., Zhang, S., Bao, F., Wang, P., Huang, X., Li, J. (2013). Bayesian network analysis reveals alterations to default mode network connectivity in individuals at risk for alzheimer’s disease. PLoS ONE, 8 (12), 1–10, 10.1371/journal.pone.0082104

Lin, A., Yang, R., Dorkenwald, S., Matsliah, A., Sterling, A.R., Schlegel, P., … et al. (2024). Network statistics of the whole-brain connectome of drosophila. Nature, 634 (8032), 153–165, 10.1038/s41586-024-07968-y

Liu, J., Ji, J., Yao, L., Zhang, A. (2019, Nov). Estimating brain effective connectivity in fmri data by non-stationary dynamic bayesian networks. 2019 IEEE International Conference on Bioinformatics and Biomedicine (BIBM), 834–839, 10.1109/bibm47256.2019.8982982

McNally, R.J., Mair, P., Mugno, B.L., Riemann, B.C. (2017). Co-morbid obsessive–compulsive disorder and depression: a bayesian network approach. Psychological Medicine, 47 (7), 1204–1214, 10.1017/S0033291716003287

Mutlu, A.Y., & Aviyente, S. (2009). Inferring effective connectivity in the brain from eeg time series using dynamic bayesian networks. 2009 annual international conference of the ieee engineering in medicine and biology society (pp. 4739–4742).

Novelli, L., & Lizier, J.T. (2021). Inferring network properties from time series using transfer entropy and mutual information: Validation of multivariate versus bivariate approaches. Network Neuroscience, 5 (2), 373–404, 10.1162/netn_a_00178

Owens, M.M., Albaugh, M.D., Allgaier, N., Yuan, D., Robert, G., Cupertino, R.B., … et al. (2022). Bayesian causal network modeling suggests adolescent cannabis use accelerates prefrontal cortical thinning. Translational Psychiatry, 12 (1), 188, 10.1038/s41398-022-01956-4

Phillips, N.S., Rao, V., Kmetz, L., Vela, R., Medick, S., Krull, K., Kesler, S.R. (2022, May). Changes in brain functional and effective connectivity after treatment for breast cancer and implications for intervention targets. Brain Connectivity, 12 (4), 385–397, 10.1089/brain.2021.0049

Pulver, S.R., Bayley, T.G., Taylor, A.L., Berni, J., Bate, M., Hedwig, B. (2015). Imaging fictive locomotor patterns in larval drosophila. Journal of Neurophysiology, 114 (5), 2564–2577, 10.1152/jn.00731.2015

Rajapakse, J., Wang, Y., Zheng, X., Zhou, J. (2008). Probabilistic framework for brain connectivity from functional mr images. IEEE Transactions on Medical Imaging, 27 (6), 825–833, 10.1109/tmi.2008.915672

Rajapakse, J.C., & Zhou, J. (2007, Sep). Learning effective brain connectivity with dynamic bayesian networks. NeuroImage, 37 (3), 749–760, 10.1016/j.neuroimage.2007.06.003

Ramakrishna, J.S., & Ramasangu, H. (2019). Estimation of functional connectivity in cognitive impaired brain using non-homogeneous dynamic bayesian model. Tencon 2019 - 2019 ieee region 10 conference (tencon) (p. 2154–2159).

Schwarz, G. (1978). Estimating the dimension of a model. The Annals of Statistics, 6 (2), 461–464, 10.1214/aos/1176344136

Scutari, M. (2010). Learning bayesian networks with the bnlearn R package. Journal of Statistical Software, 35 (3), 1–22, 10.18637/jss.v035.i03

Scutari, M., Graafland, C.E., Gutiérrez, J.M. (2019). Who learns better bayesian network structures: Accuracy and speed of structure learning algorithms. International Journal of Approximate Reasoning, 115, 235–253, 10.1016/j.ijar.2019.10.003

Seth, A.K., Barrett, A.B., Barnett, L. (2015). Granger causality analysis in neuroscience and neuroimaging. The Journal of Neuroscience, 35 (8), 3293–3297, 10.1523/jneurosci.4399-14.2015

Shojaie, A., & Fox, E.B. (2022). Granger causality: A review and recent advances. Annual Review of Statistics and Its Application, 9 (1), 289–319, 10.1146/annurev-statistics-040120-010930

Smith, V.A. (2010). Revealing structure of complex biological systems using bayesian networks. In E. Estrada, M. Fox, D.J. Higham, & G.-L. Oppo (Eds.), Network science (p. 185–204). London: Springer London.

Smith, V.A., Yu, J., Smulders, T.V., Hartemink, A.J., Jarvis, E.D. (2006). Computational inference of neural information flow networks. PLoS Computational Biology, 2 (11), e161, 10.1371/journal.pcbi.0020161

Steck, H., & Jaakkola, T.S. (2002). On the dirichlet prior and bayesian regularization. Proceedings of the 16th international conference on neural information processing systems (p. 713–720). Cambridge, MA, USA: MIT Press.

Takemura, S.-y., Hayworth, K.J., Huang, G.B., Januszewski, M., Lu, Z., Marin, E.C., … Berg, S. (2024). A connectome of the male drosophila ventral nerve cord. eLife,, 10.7554/elife.97769.1

Tsukahara, V.H.B., de Oliveira Júnior, J.N., de Oliveira Barth, V.B., de Oliveira, J.C., Rosa Cota, V., Maciel, C.D. (2022). Data-driven network dynamical model of rat brains during acute ictogenesis. Frontiers in Neural Circuits, 16, 747910, 10.3389/fncir.2022.747910

Ulusoy, I., & Geduk, S. (2024). Improved brain effective connectivity modelling by dynamic bayesian networks. Journal of Neuroscience Methods, 409, 110211, 10.1016/j.jneumeth.2024.110211

Valdes-Sosa, P.A., Roebroeck, A., Daunizeau, J., Friston, K. (2011). Effective connectivity: Influence, causality and biophysical modeling. NeuroImage, 58 (2), 339–361, 10.1016/j.neuroimage.2011.03.058

Vogels, T.P., & Abbott, L.F. (2005). Signal propagation and logic gating in networks of integrate-and-fire neurons. Journal of Neuroscience, 25 (46), 10786–10795, 10.1523/JNEUROSCI.3508-05.2005

Wang, Y.X., Li, L., Li, J.J., Huang, H. (2021). Network modeling in biology: Statistical methods for gene and brain networks. Statistical Science, 36 (1), 89–108, 10.1214/20-sts792

Warnick, R., Guindani, M., Erhardt, E., Allen, E., Calhoun, V., Vannucci, M. (2018). A bayesian approach for estimating dynamic functional network connectivity in fmri data. Journal of the American Statistical Association, 113 (521), 134–151, 10.1080/01621459.2017.1379404

Wilson, H.R., & Cowan, J.D. (1972). Excitatory and inhibitory interactions in localized populations of model neurons. Biophysical journal, 12 (1), 1–24, 10.1016/s0006-3495(72)86068-5

Winding, M., Pedigo, B.D., Barnes, C.L., Patsolic, H.G., Park, Y., Kazimiers, T., … Zlatic, M. (2023). The connectome of an insect brain. Science, 379 (6636), eadd9330, 10.1126/science.add9330

Wu, X., Wen, X., Li, J., Yao, L. (2013). A new dynamic bayesian network approach for determining effective connectivity from fmri data. Neural Computing and Applications, 24 (1), 91–97, 10.1007/s00521-013-1465-0

Xing, F., Fan, S., Dal Monte, O., Jadi, M.P., Chang, S.W., Nandy, A.S. (2025). Causal dynamics of social gaze in primate prefrontal-amygdala networks revealed by dynamic bayesian modeling. bioRxiv [Preprint],, 10.1101/2025.08.08.669405

Yu, M., Sporns, O., Saykin, A.J. (2021). The human connectome in alzheimer disease — relationship to biomarkers and genetics. Nature Reviews Neurology, 17 (9), 545–563, 10.1038/s41582-021-00529-1

Zhang, H., Benz, H.L., Bezerianos, A., Acharya, S., Crone, N.E., Maybhate, A., … Thakor, N.V. (2010). Connectivity mapping of the human ecog during a motor task with a time-varying dynamic bayesian network. 2010 annual international conference of the ieee engineering in medicine and biology (p. 130–133).

Zhang, L., Samaras, D., Alia-Klein, N., Volkow, N., Goldstein, R. (2005). Modeling neuronal interactivity using dynamic bayesian networks. Y. Weiss, B. Schölkopf, & J. Platt (Eds.), Advances in neural information processing systems (Vol. 18, p. 1593–1600). Cambridge, MA, USA: MIT Press.

